# Eukaryote-to-eukaryote gene transfer pervades the genome evolution of Rhizaria

**DOI:** 10.1101/2023.01.27.525846

**Authors:** Jolien J.E. van Hooff, Laura Eme

## Abstract

Lateral gene transfer (LGT) has contributed to the genetic makeup of various eukaryotic lineages, yet its prevalence and long-term significance remain poorly understood, particularly for transfers between eukaryotes. Here, we investigate LGT across 29 species of Rhizaria, an ancient and ecologically diverse clade of predominantly free-living, single-celled phagotrophs. Using phylogenetic analyses of over 40,000 gene families complemented by machine learning-based prediction, we estimate that 8–20% of protein-coding genes in contemporary rhizarian genomes were acquired through LGT at various points during their billion-year history, with ~2,000 transfer events shared between at least two species across the rhizarian tree of life. Gene duplications outnumber LGTs across most lineages, yet LGT-derived genes themselves duplicate more frequently than vertically inherited ones, amplifying the genomic impact of each transfer event. Notably, transfers from other eukaryotes outnumber those from prokaryotes and show distinct signatures: prokaryote-derived LGTs are enriched among extracellular proteins, whereas eukaryote-derived LGTs are overrepresented in nuclear and informational processes. Prokaryote-derived LGT genes progressively acquire introns over evolutionary time, confirming their genomic integration and long-term retention. Our findings establish LGT as a pervasive force in rhizarian genome evolution and highlight eukaryote-to-eukaryote transfer as a substantial but often overlooked component of eukaryotic genetic innovation.

## Introduction

Lateral gene transfer (LGT) refers to the non-inheritance transfer of genetic material between unrelated species, and is well-documented among bacteria and archaea. Its role and prevalence in eukaryotes have been more controversial and less clear, although increasing evidence suggests that LGT has played a significant role in the evolution of many eukaryotic lineages^1–4^. Questions persist around the molecular and cellular mechanisms underlying these transfers, their relative importance compared to gene duplication, and their long-term effects on the recipient genomes^5–7^. Myriad studies indicate that LGT has helped eukaryotes acquire new capacities, such as establishing stable relationships with their endosymbionts, adapting to or reverting from parasitism, or thriving in low-oxygen environments^8–12^. Recently, LGT-derived genes were estimated to constitute 1% of the total gene inventories of algae and protists^5^, which hints at a significant, but modest role of LGT in eukaryotic evolution.

Several factors complicate our understanding of LGT in eukaryotes. Primarily, research has often been directed at pinpointing LGTs that are unique to specific organisms, focusing largely on recent events. This significantly limits our ability to gauge the long-term significance of LGT and hinders a consistent comparison of LGT frequencies across different organisms using a standard set of criteria. Consequently, it remains unclear how the frequency of LGT compares to other evolutionary processes for gene acquisition, such as gene duplication or *de novo* gene invention. Additionally, the study of LGT between eukaryotic organisms has been relatively neglected compared to LGT from prokaryotes to eukaryotes, largely due to the inherent challenges in detection. State-of-the-art methodologies, which rely on discrepancies between gene and species trees, often fall short when analyzing closely related lineages. This is because single gene or protein phylogenies, derived from a limited phylogenetic signal in a small number of analyzed sites, may not be sufficiently resolved. In addition, independent gene losses across the tree can erroneously suggest LGT events, by grouping sequences from “distant” species. As a result, single gene phylogenies frequently lack the necessary information to differentiate LGT between eukaryotes from vertical gene transmission confidently. On top of that, we still lack high-quality genome data for many eukaryotes^13–15^, especially for microbial eukaryotes, while such data underpin the most sophisticated evolutionary analyses, including the detection of LGTs with phylogenetics.

The Rhizaria represent an interesting, largely unexplored, lineage for addressing these evolutionary questions. This billion-year old clade of predominantly single-celled, free-living phagotrophs exhibits substantial morphological and ecological diversity, reflecting adaptation across varied habitats. Despite their evolutionary success and the complex feeding behaviors that may predispose them to genetic exchange, Rhizaria remain genomically understudied^15,16^. Many of its groups are represented primarily by transcriptomic data or fragmented genome assemblies, which has hindered large-scale phylogenomic investigations.

In this work, we investigated the contribution of LGT to Rhizaria genomes by identifying and characterizing genes acquired from both prokaryotic and eukaryotic donors, and by evaluating the roles of LGT and gene duplication in shaping genome dynamics. We detected LGT events throughout the Rhizaria tree of life, including ancient transfers, which allowed us to characterize the evolutionary trajectories of transferred genes, and revealed that they undergo frequent duplication and loss. We further compared LGT-derived proteins with vertically inherited ones, then leveraged their distinguishing features to train a machine learning model for predicting LGT-derived proteins without phylogenies. In summary, our study offers a detailed quantitative and qualitative exploration of LGT within the largely unexplored Rhizaria lineage, emphasizing the significant impact of foreign genes on the genetic makeup of this predominantly microbial group.

## Results

### Extensive gene acquisition in Rhizaria through LGT from prokaryotes and eukaryotes

To investigate LGTs in Rhizaria, we aimed to develop a sophisticated phylogeny-based approach, attuned to the data-quality shortcomings in this poorly studied group. We first collected protein sequences from 29 rhizarians, most of which (27) were derived from transcriptome or low-quality genome data, with some being relatively incomplete, as BUSCO scores indicate (Figure 1, Supplementary Table 1). We collected their homologs from other eukaryotes, prokaryotes and viruses. After discarding protein families displaying potentially spurious or weak signals (Methods), we inferred phylogenies for 40,951 families. In each tree, we identified the monophyletic groups of rhizarian sequences (we will also refer to these as ‘rhizarian clades’) and determined their evolutionary origin. Briefly, after removing potential contaminants based on suspiciously high sequence identities, and for genomes, validating contig authenticity (Methods), and taking into account statistical support, we designated each rhizarian clade as 1) having a vertical origin, if it forms a monophyletic group with stramenopiles or alveolates (the closest relatives of rhizarians, all together forming the SAR group^17–19^), 2), having a lateral origin, if it is nested within a specific non-SAR clade supported by at least two well-resolved phylogenetic nodes, or 3), representing a gene invention, if the tree only contains rhizarian sequences. We also conducted gene tree-species tree reconciliation of the subtrees embodied by the monophyletic rhizarian groups, in order to chart gene duplication and loss events within Rhizaria. For this, we inferred a Rhizaria species phylogeny, and subsequently mapped the LGTs, gene duplications and inventions onto it (Methods, Figure 1).

**Figure 1.**
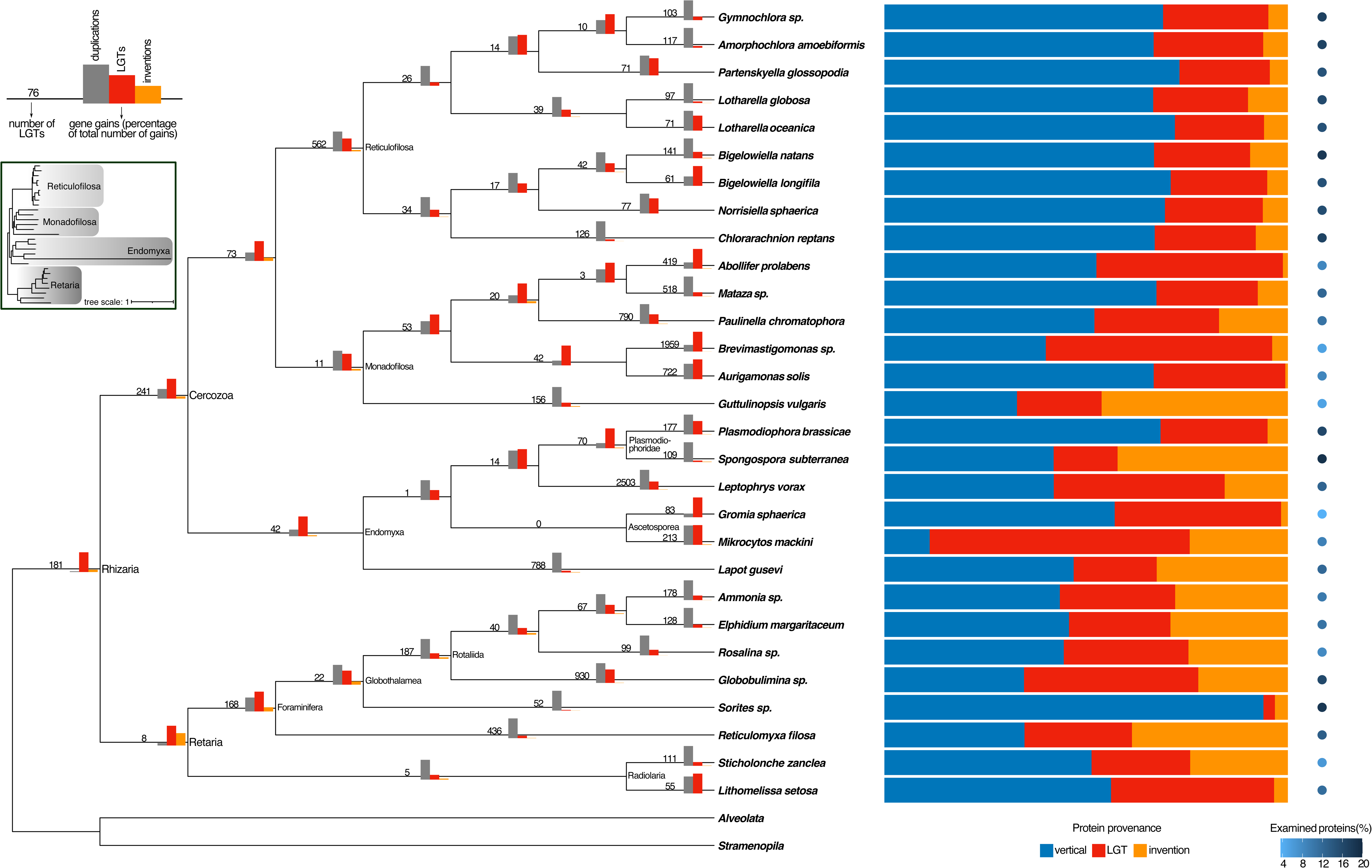
LGT, duplication and invention events across the rhizarian tree of life. Absolute LGT event counts are displayed on each branch, while bar charts atop the branches show the relative frequencies of LGT, duplication, and invention events. Adjacent to the tree, stacked bar charts illustrate the contributions of vertical inheritance, LGT, and gene invention to the gene inventories of each existing species. A blue gradient dot to the right signifies the proportion of proteins analyzed through our phylogenetic process relative to the total protein count. An inset details the Rhizarian phylogeny with accurate branch lengths, with the species placed in the same order as the schematic tree, and displays the four major clades within Rhizaria. The scale bar denotes the expected number of substitutions per site.

#### Phylogeny-based LGT inferences: maximum vs shared estimates

We detected 13,282 LGT events across the Rhizaria tree of life (Figure 1). Interpreting their present-day imprint requires specifying the dataset and denominator used (Table 1). Under our phylogeny-based dataset—i.e., the subset of proteins that passed all tree-building and contamination filters and for which an origin could be assigned phylogenetically (12% of all proteins in the 29 proteomes; Table 1)—LGT-derived proteins account for an average of 30% of the analyzed proteins across species (Table 1; Supplementary Table 1). We treat this value as an upper bound because this dataset includes many species-specific rhizarian clades, which are intrinsically hard to validate in transcriptome-derived proteomes. Noticeably, the good-quality genomes, for which we validated that candidate LGT genes are located on contaminant-free contigs (Methods), display comparable LGT percentages (*Bigelowiella natans*: 24% and *Plasmodiophora brassicae*: 27%). Based on this, we consider our LGT account of transcriptomes to be plausible, despite eventual residual contamination.

**Table 1.** Summary of LGT quantification metrics and respective datasets.

| Estimate category | Value (%) | Numerator | Denominator | Context |
| --- | --- | --- | --- | --- |
| **Phylogenetically analyzed proteome - <u>maximum</u>** | **12%** | Proteins inferred an origin type based on phylogenetic analysis of 40,951 gene families: 99,974 | Total proteins across 29 rhizarian species: 835,552 | The subset of proteins passing all quality filters for tree-building and subsequent tree analysis, including contamination filters.<br>Figure 1, Supplementary Table 1 (2nd sheet) |
| **Average LGT percentage across species - <u>maximum</u>** | **30%** | Proteins with LGT origin: 28,298 in total for all species combined - NB: The average percentage across the 29 species is used. | Proteins analyzed via the maximum phylogenetic approach: 99,974 in total for all 29 species | The upper bound of the LGT signal detected within the analyzed subset.<br>LGT origin proteins stem from the 13,282 inferred LGT events.<br>Figure 1, Supplementary Table 1 (2nd sheet) |
| **Phylogenetically analyzed proteome - <u>shared</u>** | **6%** | Proteins inferred an origin type based on phylogenetic analysis of 40,951 gene families, including only multi-species clades: 49,266 | Total proteins across 29 rhizarian species: 835,552 | The subset of proteins passing all quality filters for tree-building and subsequent tree analysis, including contamination filters, with orthologs in other rhizarians of the dataset.<br>Supplementary Figure 1, Supplementary Table 1 |
| **Average LGT percentage across species - <u>shared</u>** | **20%** | Proteins with LGT origin based on shared set: 10,461 in total for all species - NB: The average percentage across the 29 species is used. | Proteins analyzed via phylogeny under the shared constraint: 49,266 in total for all 29 species | Highlights the long-term impact of LGT while excluding potential contamination.<br>LGT origin proteins stem from 1,992 shared (non-species-specific) LGT events<br>Supplementary Figure 1, Supplementary Table 1 |
| **Analyzed proteome - <u>combined</u> (maximum + ML)** | **72%** | Proteins inferred an origin type based on the combined approach: 601,199 | Total proteins across 29 rhizarian species: 835,552 | The subset of proteins passing all quality filters for tree-building and subsequent tree analysis, including contamination filters, and those that do not, but are subjected to occurrence and ML-based inference (Methods)<br>Supplementary Figure 2, Supplementary Table 1 |
| **Average LGT percentage across species - <u>combined</u>** | **8%** | Proteins with LGT origin based on combined approach: 48,139 in total for all species - NB: The average percentage across the 29 species is used. | Proteins analyzed via combined approach: 601,199 in total for all 29 species | Our most global, conservative lower bound estimate.<br>Supplementary Figure 2, Supplementary Table 1 |

To derive a conservative estimate that is robust to contamination concerns in transcriptome datasets, we repeated the phylogeny-based analysis while excluding all rhizarian clades represented by a single species (a “shared” constraint). This yields 1,992 shared (non–species-specific) LGT events distributed across internal branches of the rhizarian tree, which account for ~20% of proteins within the shared phylogenetic dataset (6% of all proteins across the 29 proteomes; Table 1; Supplementary Figure 1; Supplementary Table 1). Notably, a substantial fraction of this shared signal maps to deep rhizarian ancestors (e.g., Reticulofilosa, Monadofilosa, Rotaliida), indicating long-term retention of transferred genes (Figure 1; Supplementary Figure 1).

#### Prediction-assisted proteome-wide LGT prevalence: a conservative lower-bound estimate

Because only a minority of proteins pass stringent phylogeny-based filtering (12% in the maximum dataset; 6% in the shared dataset; Table 1), we additionally estimated a conservative genome-wide lower bound for LGT using a non-phylogenetic workflow that combines lineage distribution–based labeling with a Gradient Boosting classifier trained on phylogeny-derived calls (Methods; Supplementary Text). This combined approach assigns an origin to 72% of proteins across the 29 proteomes. It infers that, on average, ~8% of analyzed proteins originated via LGT (Table 1; Supplementary Figure 2; Supplementary Table 1). Because the classifier has a non-negligible false-positive rate and does not provide gene-by-gene phylogenetic validation, we use this estimate only to contextualize the global scale of LGT in Rhizaria; all downstream biological characterization (Figures 2–4) is restricted to the phylogeny-based LGT set unless stated otherwise.

**Figure 2.**
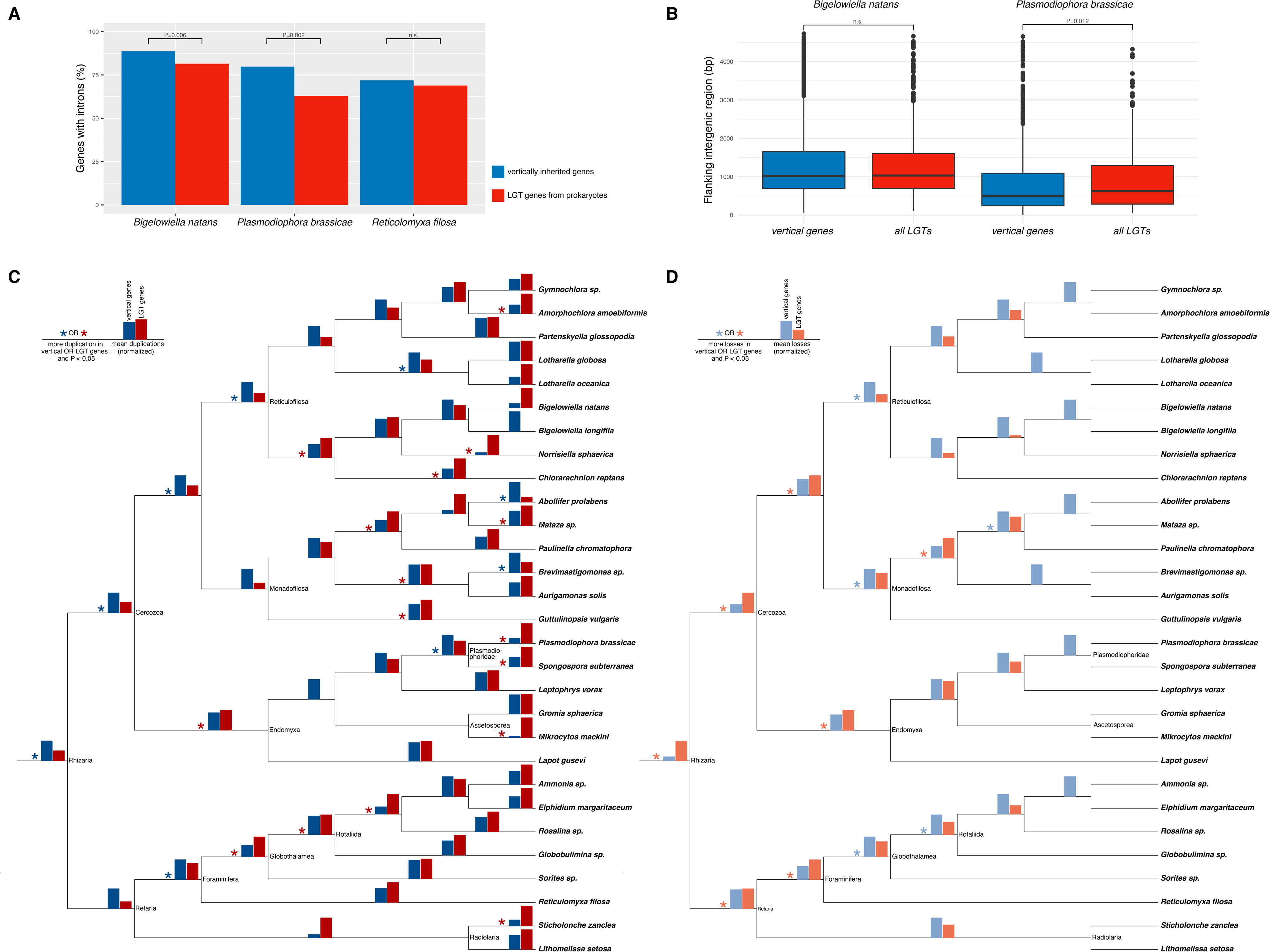
Evolutionary fates and genomic contexts of genes acquired by LGT. **A**. Fraction of vertically inherited and LGT-derived genes displaying at least one intron. Intron presence was only assessed for three species with appropriate genomic data: *B. natans*, *P. brassicae* and *R. filosa*. P-values were obtained with the chi-square test of independence. **B**. Distributions of the flanking intergenic region (FIR) lengths for LGTs and vertically inherited genes in *B. natans* and *P. brassicae*. P-values were obtained with the Mann-Whitney U-test. **C,D**. Gene duplications (C) and losses (D) (mean of normalized values, see Methods) for vertically inherited genes and genes acquired by LGT across lineages in the rhizarian tree of life. Note that no LGT was inferred at the base of ‘Ascetosporea’ (Figure 1). Losses could only be inferred for non-terminal branches. P-values were obtained with the Mann-Whitney U-test.

**Figure 3.**
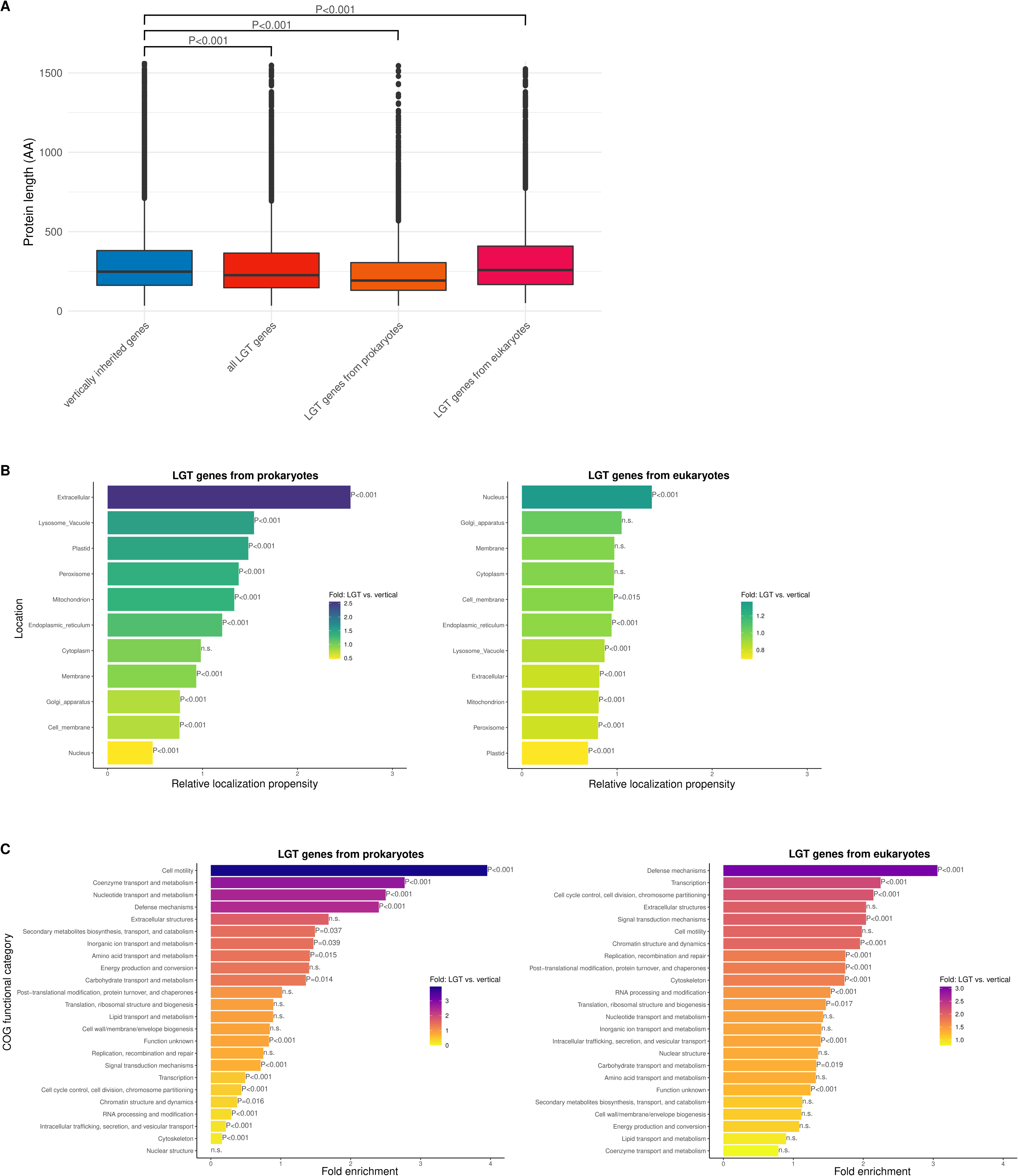
Features of proteins acquired by LGT. **A**. Distributions of protein lengths of vertically inherited genes, all proteins acquired by LGT, proteins acquired by LGT from prokaryotes only, and from eukaryotes only. P-values were derived from a Mann-Whitney U test on the distributions of the protein lengths for LGT-derived versus vertically inherited proteins. The protein lengths used as input are supposed to approximate the length upon transfer, or vertical inheritance into eukaryotes, by taking the median of all rhizarian sequences in each ancient rhizarian clade in the gene phylogenies. **B**. Localization propensities of LGT proteins compared to vertically inherited proteins based on DeepLoc predictions and subsequent projection onto ancestral nodes in single gene trees, reflecting the actual LGT, or vertically inherited, protein (Methods). If relative localization propensity >1, the LGTs have on average a stronger predicted probability to localize to a specific cellular compartment than vertically inherited genes. LGTs are divided based on prokaryotic or eukaryotic donor sources. Significance of the differences between vertically inherited and LGT proteins was assessed with a Mann-Whitney U test on the distributions of the localization probabilities. **C**. Fold-enrichment of LGT proteins in COG functional categories compared to vertically inherited genes, split by donor. Like for (A) and (B), this analysis considers the features, in this case, the predicted annotations, at the node uniting the ancestral rhizarian clade in single gene trees (Methods). Chi-square tests assessed the significance of differences in the proportion of LGT and vertically inherited proteins within each category.

**Figure 4.**
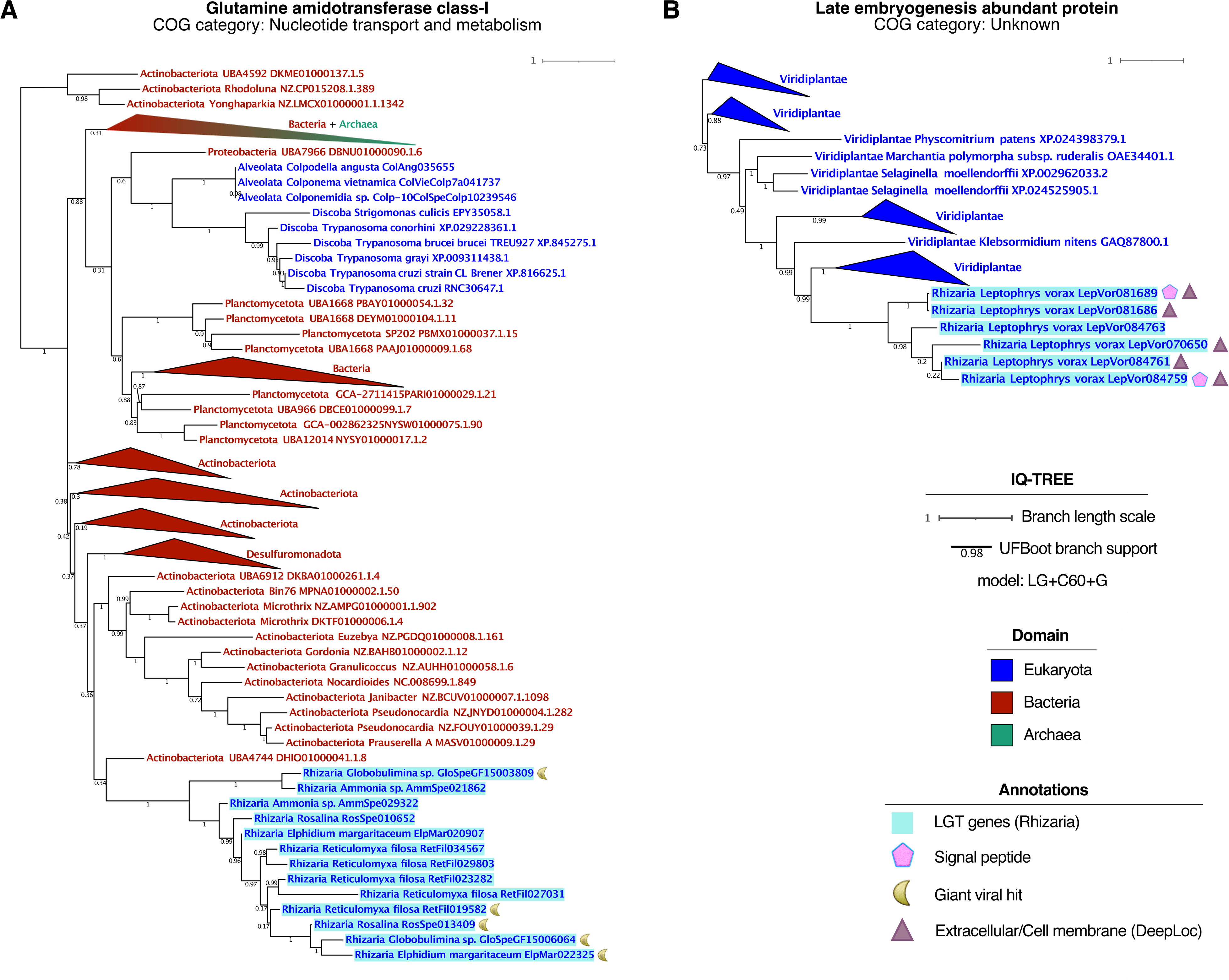
LGT cases illustrating some of the main patterns identified across Rhizaria. **A-B**. Maximum-likelihood phylogenies inferred for selected LGTs with IQ-TREE under the LG+C60+G evolutionary model. The rhizarian sequences descending from LGT events are highlighted in light blue. The description comprises the most informative annotation from the rhizarian sequences themselves or from their homologs in other taxa. The ‘COG category’ summarizes the EggNOG-mapper annotations of all sequences descending from the LGT. Signal peptides were predicted with TargetP and cell membrane/extracellular localization of proteins with DeepLoc. Putative homologs in giant viruses were identified using hmmscan of the rhizarian proteins against HMM profiles from GVOG.

#### Temporal context from genome lineages

To place the magnitude of detectable LGT into temporal context, we computed a “lineage-specific retained-event accumulation metric” for the two rhizarians with high-quality genomes (*P. brassicae* and *B. natans)*. Specifically, for each focal lineage we summed the LGT events mapped between the last common ancestor of Rhizaria and the terminal branch leading to the focal species (Figure 1). For *Plasmodiophora brassicae*, we infer 726 LGT events along this lineage, corresponding to ~0.37–0.43 retained detectable events per million years using SAR divergence time estimates of 1,673–1,986 Mya^20^ (Figure 1). For *Bigelowiella natans*, the analogous value is ~0.65–0.77 retained detectable events per million years (Figure 1). Including additional events inferred by the combined phylogeny+prediction approach increases these lineage-specific values (Supplementary Figure 2), but we emphasize that all such estimates reflect present-day detectability and therefore conflate acquisition with subsequent gene loss, incomplete taxon sampling, and detection limits. Accordingly, they should be interpreted as lower bounds on historical LGT activity, rather than direct estimates of underlying transfer rates.

#### LGT versus gene duplication across the rhizarian tree

We then aimed to assess the visible imprint of LGT onto modern genomes, compared to gene duplications (Figure 1) – the mechanism generally assumed to be the main source of genomic novelty in eukaryotes. Comparing the total numbers LGT and duplication events in the entire Rhizaria tree of life, the latter occurred three times more frequently. Looking at individual branches, we observed that 36 of the 57 lineages underwent more duplications than LGTs. Notable exceptions included the ancestors of Rhizaria (181 LGTs vs. 7 duplications) and Cercozoa (241 LGTs vs. 117 duplications). The prevalence of LGT compared to duplications in different branches seems to be affected by the clade they are part of. For instance, Monadofilosea branches mostly experienced more LGTs than duplications, whereas the opposite was true for Foraminifera. The high duplication rate in foraminiferans might be attributed to their frequent endoreplication processes^21^, which could lead to gene duplications as a side effect. These processes might also necessitate the separation of germline from somatic genomes^21^, potentially making LGT fixation more challenging, especially if foreign genes are inserted into somatic rather than germline genome. Notably, like studying LGTs, studying gene duplications remains very challenging if one is limited to (mostly) lower-quality genomes and transcriptomes as primary data source, because they might artificially inflate (e.g., due to splice variants) or deflate (e.g., due to gaps in the assemblies) duplicate counts. Yet, the higher numbers of duplication events, relative to LGT events, apply to the branches with the better-quality genomes in our dataset too (*B. natans*, *P. brassicae)*, and therefore we deem this a solidly discerned pattern.

#### Eukaryote-derived transfers dominate

We determined the source of each LGT by identifying the clade within which the Rhizaria are nested. We found that 48% of the LGTs originated from non-SAR eukaryotes, 39% from bacteria, and less than 1% from archaea. The remaining LGTs were of unclear origin. Most individual branches (46 out of 57) received more LGTs from eukaryotic sources compared to prokaryotic ones (Supplementary Figure 3, Supplementary Table 4). The exceptions were observed at the tips, rather than deeper within the tree, either indicating residual contamination in the sequence data or suggesting that eukaryote-derived genes are more likely to be preserved over the long term than those from prokaryotes. Hence, when excluding LGTs limited to a single species (Supplementary Figure 1), the majority was eukaryotic: 51%, in contrast to 26% and less than 1% from bacteria and archaea, respectively. Many branches seem enriched for LGTs from Opisthokonta, such as Foraminifera, for which 26% of all LGTs seemed of opisthokont origin (Supplementary Table 1). This Opisthokonta dominance might result from bias in the composition of the database, which is a limiting factor for pinpointing donors. Yet, the ancestors of Foraminifera, and possibly also early Cercozoa, might have adopted eukaryvorous feeding, which may have exposed them to many eukaryotic stretches of DNA. Intriguingly, this feeding behaviour might also have impacted their cell sizes^22^. Possibly, influxes of eukaryotic genes were not only a consequence of eukaryovory, but also a cause, as these genes might have effectuated eukaryovory-facilitating features, like the ability to grow larger. *Paulinella chromatophora*, a unique species that acquired a primary photosynthetic organelle independently of Archaeplastida, had many bacterial LGTs (402/790, 51%), as reported before^23^, yet - and this, to our knowledge, has not been observed yet - it also received many eukaryotic LGTs (299 out of 790, 38%).

### The fate of laterally transferred genes

#### Integration via intron acquisition

We sought to determine how frequently genes of prokaryotic origin assimilated to the host genome through spliceosomal intron acquisition^12^. In the three species for which we have adequate genomic data, most prokaryote-derived LGT genes contained at least one intron (Figure 2A, *B. natans*: 81% of prokaryote-derived LGT genes, *P. brassicae*: 63%, *Reticulomyxa filosa*: 69%). This confirms that these prokaryote-origin genes are not contaminants. Moreover, the canonical lengths of these introns (Supplementary Figure 4B) suggest that these genes are expressed by the recipient, a hypothesis that awaits validation from transcriptomic data. Noticeably, the youngest, lineage-specific LGT genes lack introns more often than older LGTs (Supplementary Figure 4A), suggesting a gradual intron acquisition over time (for more details, see Supplementary Text: ‘Introns in prokaryotic-derived LGTs’).

Our intron-based analyses are most informative for validating genomic integration of prokaryote-derived LGT candidates in the few rhizarians with genome assemblies. By contrast, eukaryote-to-eukaryote LGT could occur either via DNA transfer or via an RNA intermediate followed by reverse transcription and integration, in which case transferred genes would be expected to be intron-poor and could resemble retrogene-like insertions. Because our intron analyses were restricted to genome assemblies and primarily used to support integration of prokaryote-derived candidates, we do not infer transfer mechanisms for eukaryote-derived LGT from intron presence/absence. Disentangling DNA- versus RNA-mediated routes for eukaryotic LGT will require broader genome sampling and targeted analyses of gene structure and integration signatures.

#### Post-transfer dynamics: duplication and loss relative to native genes

To gauge the impact of LGTs on the host organisms, we examined their evolutionary trajectories after transfer. If LGT genes duplicate frequently, they might confer a beneficial function, possibly benefiting from increased gene dosage. Although duplications of LGT genes have been documented^10,11,24–28^, a systematic comparison with native genes is pending. Here, we found that in most lineages (38/56), LGT genes duplicated more often than vertically inherited genes (Figure 2C), with significant differences observed in 16 lineages. This suggests these genes not only fulfill crucial adaptive roles, but also that their duplications contribute to the large proportion of LGT-derived genes in modern-day rhizarian genomes. We also examined whether LGT genes were lost more frequently than vertically inherited genes, as this would indicate their relatively transient impact on lineages. In the limited lineages with statistically significant differences, seven showed a higher loss rate for LGT genes, compared to five for vertically inherited genes, showing no distinct loss tendencies (Figure 2D).

#### Molecular remodeling after transfer: sequence divergence and domain turnover

Beyond duplication and loss, we explored the sequence evolution rates of LGTs and their propensity for protein domain gain and loss. Our findings indicate that LGT proteins evolve significantly faster than vertically inherited genes in certain lineages (Supplementary Text, ‘LGT sequence divergence’). Moreover, they tend to gain and lose domains more frequently across most lineages (Supplementary Text, ‘Domain loss and gain’). In conclusion, our analysis suggests that LGT genes evolve more dynamically than vertically inherited genes.

### Mechanisms and constraints shaping transfer and integration

#### Viral footprints in LGT phylogenies

Given that viruses are known to facilitate LGT in eukaryotes^29^, we searched all gene trees for viral sequences. We found that in our phylogenies, clusters of rhizarian proteins originating from LGT were more likely to contain viral sequences than those inherited vertically (18% *versus* 13%, respectively, P<0.001). Although this finding does not provide direct evidence, it indicates viral–eukaryotic gene transfer to occur frequently, regardless of directionality^30^, and suggests that viruses could have played a role in mediating some LGT events.

#### Insertion landscape in genomes: gene density and intragenomic relocation

We also investigated the genomic context of laterally acquired genes. LGTs might preferentially localize to gene-sparse, potentially heterochromatic areas of the genome to avoid disrupting the genetic integrity of the host^12^. We examined the gene density surrounding LGTs in *B. natans* and *P. brassicae* by analyzing the flanking intergenic regions (FIRs), representing the distances to neighboring genes. For *B. natans*, we observed no notable difference in FIR lengths between LGTs and vertically inherited genes (Figure 2B). In contrast, in *P. brassicae*, LGTs were found to have significantly larger FIRs compared to vertically inherited genes (median FIR for LGTs: 628 bp, median FIR for vertically inherited genes: 505 bp, P=0.012), indicating a preference for integration into gene-sparse areas. Furthermore, we investigated whether LGTs initially integrated into sparse genomic regions and subsequently relocated to denser, potentially more transcriptionally active areas, as suggested^12^. Our comparison of FIR lengths for LGTs acquired at various time points revealed that, in *P. brassicae*, the most recent, species-specific LGTs are situated in the sparsest genomic regions (median FIR: 777 bp, Supplementary Figure 4E). This pattern could suggest that LGTs indeed initially insert in areas where they cause the least harm and possibly are transcriptionally repressed.

#### Structural constraints on transfer: protein length patterns depend on donor

Finally, to shed light on putative structural constraints on transfer, we comprehensively analyzed structural features of LGT and non-LGT proteins. Several studies reported that LGT genes are typically short^24,31,32^, hypothetically because shorter genes are more likely to be fully integrated and functional upon transfer, even if only a small fragment of foreign DNA is incorporated. Our analysis indeed shows that LGT proteins are generally slightly shorter than their native counterparts (Figure 3A, median protein lengths: 226 (LGT proteins) *versus* 248 (native proteins) amino acids (P<0.001). However, proteins from eukaryotic LGTs are slightly longer than native proteins (median length: 258, P<0.001), hence the observed shortness specifically pertains to prokaryotic LGTs (median length: 192). This difference likely reflects the intrinsic length disparity between eukaryotic and prokaryotic proteins^33^ (also see Supplementary Text: ‘Protein lengths across datasets’). In general, the lengths of LGT and non-LGT proteins differ only marginally, so we cannot conclude that the integration of larger DNA segments poses a major barrier to LGT.

### Transferred proteins harbor donor-specific signatures

#### Subcellular localization biases differ for prokaryotic vs eukaryotic donors

To explore the roles LGT genes play within their new hosts, we employed DeepLoc^34^ to predict proteins’ localizations and EggNOG-mapper^35^ to functionally annotate them. Similar to protein length, the signatures of LGTs differ between those donated by prokaryotes and eukaryotes. Moreover, the predicted localizations of genes of prokaryotic origin deviate more strongly from those of native genes, than those of eukaryotic origin. Prokaryotic LGTs are predicted to localize extracellularly 2.6 times more frequently than vertically-inherited genes (P<0.001; Figure 3B, 803 LGTs in total; Supplementary Figure 5E), suggesting a role in host-environment interactions. Such interactions could comprise species-species interactions and public goods cooperation, as suggested for bacterial secreted LGT proteins^36^, as well as oomycete LGTs^37^. LGTs’ potential roles in interspecies interactions is suggested by significant enrichments in functional categories such as ‘Defense mechanisms’ for prokaryotic LGTs (2.4-fold enrichment, respectively; P<0.001; Figure 3C), despite low absolute LGT numbers in this category (36, respectively, Supplementary Figure 5G).

Eukaryotic LGTs are most enriched in the nucleus (1.4-fold higher average localization probability compared to vertically inherited genes, P<0.001; Figure 3B, 1515 LGTs in total; Supplementary Figure 5F), a surprising and significant finding without an immediate explanation in terms of adaptive function. Interestingly, like prokaryotic ones, eukaryotic LGTs show strong overrepresentation in ‘Defense mechanisms’ (3.1-fold enrichment, P<0.001, 53 LGTs in total; Supplementary Figure 4H), forming one of the few similarities between the donor groups. More surprisingly, informational processes like ‘Transcription’ and ‘Cell cycle control’ were enriched (2.2 and 2.1-fold enrichment, P<0.001; Supplementary Figure 5H), aligning with the enriched predicted nuclear localization. This suggests that LGTs can significantly influence key informational processes, challenging previous characterizations that emphasized metabolic functions, which may have stemmed from analyzing predominantly prokaryote-to-eukaryote transfers.

#### The shared LGT subset recapitulates key signals

In addition to analyzing all phylogenetically detected LGTs (Figure 1), we examined the functional signatures of the shared LGTs only (Supplementary Figure 1), which comprise the most trustworthy subset. This analysis showed similar results, except that the prokaryote-derived LGTs were slightly less overrepresented in the extracellular environment, compared to the entire set of LGTs, and that many COG functional categories yielded insignificant results due to the smaller LGT numbers (Supplementary Figure 5I,J).

#### LGT affecting transcription and epigenetic control

We highlight the remodeling of essential processes by foreign-origin genes, such as transcription factors and epigenetic regulators. For instance, several rhizarian lineages have acquired genes encoding chromo domains, associated with histone modification and transcription regulation, from Opisthokonta (Supplementary Table 2). Similarly, Reticulofilosa acquired a translational regulator, Impact, from red algae, which might play a role in stress response translation. These findings underscore the multifaceted contributions of LGTs to the evolution and functional diversification of recipient lineages.

### Highlighted cases of LGT

We highlight two cases that exemplify recurrent features of LGT-derived genes across Rhizaria.

The first involves a Glutamine amidotransferase class-I enzyme acquired by Foraminifera from bacteria (Figure 4A), consistent with the enrichment of prokaryotic LGTs in the COG category ‘Nucleotide metabolism and transport’ observed in this clade. The phylogeny suggests that homologs of this gene were independently transferred to other eukaryotic groups, including alveolates and Kinetoplastida, with possible subsequent inter-eukaryotic transfers. Within Rhizaria, and particularly in *R. filosa,*–for which a genome assembly is available, lending confidence that these are genuine paralogs–, the gene has undergone multiple post-transfer duplications, illustrating the duplication-prone dynamics of LGT genes documented above (Figure 2C). Some sequences also resemble giant viral genes (GVOGs), raising the possibility that a viral vector mediated the original transfer.

The second case involves the protoplast feeder *Leptophrys vorax*, which carries numerous LGTs predicted to encode secreted or membrane-associated proteins. The example in Figure 4B shows a family of proteins presumptively acquired from plants, annotated as ‘Late embryogenesis abundant protein 2’ (LEA-2, PF03168). Although LEA-2 function is best characterized in land plants, homologs occur in streptophyte algae and prokaryotes, where they are hypothesized to confer stress tolerance^38^. In *L. vorax*, this gene has duplicated extensively post-transfer, with most descendants retaining a predicted secretion signal or membrane localization (Figure 4B). This case thus illustrates two key patterns simultaneously: the frequent post-transfer duplication of LGT genes and the enrichment of secreted proteins among eukaryote-derived transfers.

## Discussion

Across Rhizaria, our phylogeny-based inference reveals pervasive LGT, including many events mapped to internal branches that have been retained over long evolutionary timescales. Focusing on the contamination-robust “shared” set, we infer 1,992 LGT events distributed across deep and shallow rhizarian lineages, accounting for ~20% of proteins in the shared phylogenetic dataset. This shared signal includes both prokaryote-and eukaryote-derived transfers and indicates that foreign genes have repeatedly contributed to rhizarian gene repertoires. The higher 30% estimate arises from the maximum phylogenetic dataset, which includes many species-specific clades and therefore represents an upper bound—especially in transcriptome-derived proteomes where genomic-context validation is impossible. Accordingly, we base many biological interpretations on the shared/internal-branch signal and on genome-supported analyses (e.g., introns and genomic context in *B. natans* and *P. brassicae*), while presenting broader estimates (including the 8% combined approach) as contextual bounds on the plausible contribution of LGT (Table 1). These high percentages underscore the scope and depth of our analyses, including old and eukaryote-derived LGTs, a component often overlooked.

Our findings emphasize the importance of considering LGT as a key evolutionary force shaping eukaryotic genomes. Although gene duplications are more common, LGTs represent a crucial source of genetic novelty, evidenced by the identification of over 13,000 phylogenetically detected LGT events across the Rhizarian tree of life. Ignoring LGT risks overestimating the number of species-specific genes and misunderstanding the evolutionary histories of eukaryotic genes and lineages. Moreover, LGT challenges conventional interpretations of patchy gene distributions in eukaryotes. Instead, it suggests a more dynamic evolutionary landscape than previously thought, influenced by LGT rather than mere vertical inheritance from a Last Eukaryotic Common Ancestor with an inconceivably large gene repertoire, followed by myriad losses.

Working with transcriptomic data comes with several challenges. One of them is potential contamination by foreign sequences, which might be mistaken for LGTs, specifically for organisms that cannot be grown in pure culture. While we strived to filter it out (Methods), our results likely are not completely devoid of sequence contamination. Of note, the very same factors that make Rhizaria susceptible to contamination –complex feeding behaviors and (endo)symbiotic relationships– may promote the integration of foreign genes, leading to many LGTs. This process of symbiosis-mediated LGT could, in turn, enhance the symbiotic relationships themselves, a phenomenon observed in other evolutionary lineages^39^.

In addition to filtering contaminants based on sequence identity, we rigorously discounted all species-specific clades, which suggested that 20% of the phylogenetically analyzed proteins originated from LGT (Supplementary Figure 1). This estimate represents a lower bound because it excludes species-specific LGTs, which are likely numerous (Supplementary Figure 6), also given the considerable phylogenetic distances among many species in our dataset, which in turn is due to the relatively sparse sampling of this group.

Other challenges in working with transcriptomic data pertain to their relative incompleteness, as indicated by low BUSCO scores (Supplementary Table 1), and to paralogues potentially being missed, as in many instances only one paralog is expressed. Like contamination, these risks might paint an inadequate picture of LGT and of genome evolutionary processes in general, for example by erroneously timing LGT events, overestimating losses and underestimating duplications. Hence, to fully appreciate the relative and absolute impact of LGT in this group, and to understand their patterns of genome evolution, a better (i.e., genomic), deeper and more extensive sampling of the Rhizaria clade is necessary^13,16^.

Our findings highlight the frequency and characteristics of eukaryote-to-eukaryote LGTs, distinguishing them from prokaryote-to-eukaryote ones. The latter were the subject of vastly more studies, yet eukaryote-donated genes did play important roles in the evolution of several lineages, such as transfers between closely related, cheese-adapted fungi^40^, fungal-insect pathogens^41^, fungi-to-oomycete transfers^42^ and plant-to-insect transfers^39^. A pivotal question centers on if and how these integrate into the recipient’s cellular machinery. This question particularly applies to proteins acting in nuclear and cell cycle processes, given that these involve numerous proteins, with which such an LGT might need to interact. Eukaryote-derived LGTs, however, may offer advantages for integration. For instance, the presence of introns in these genes might reduce their susceptibility to gene silencing machineries like the HUSH complex^43^. LGT may occur more readily between closely related species, as evidenced in grasses^44,45^, suggesting a propensity for transfers within similar biological contexts. This potential bias for LGTs among closely related lineages may imply that our current LGT estimates are conservative, since we did not consider LGTs from close relatives such as stramenopiles, alveolates, or other rhizarians.

How, and to what extent, LGT-derived genes and the proteins they encode differ from vertically inherited genes forms valuable patterns that machine learning-based LGT prediction might profit from. Our early exploration of such an approach indicates that features like protein length and predicted subcellular localization might inform on the likelihood that a gene was acquired through LGT, as suggested by the feature importances of our classifier (Supplementary Text, ‘Combined phylogenetics and machine learning approach estimates 8% of proteins were derived from LGT’). As prokaryote-derived proteins display more strongly distinct properties, their true origin is more likely to be accurately predictable (see Methods). Our current classifier still has a substantial false positive rate (26%, Methods), and we foresee that adding features related to the genes, based on high-quality genome sequencing, might not only help to train more accurate classifiers, but might potentially also surface indications on mechanisms involved in actual transfer and integration, like proximity to viral and TE sequences.

Importantly, our survey of LGTs within the “deep” parts of the Rhizaria phylogeny enabled us to map the trajectory of gene transfers over time, shedding light on how these genes evolved and potentially adapted within their new host. Notably, we found that older prokaryote-derived LGT genes more frequently have introns than their more recently acquired counterparts. In *P. brassicae*, we also observed that older LGT genes are situated in genomic regions with higher gene density compared to newer LGTs, suggesting that these genes migrate from areas of lower to higher gene density over time. This intragenomic migration, coupled with intron acquisition, could potentially facilitate increased gene expression. Consequently, these observations raise the question whether older LGT genes are more active than newer ones, as one might hypothesize that time plays a crucial role in the multidimensional integration of LGTs. In conclusion, our investigation into LGT in Rhizaria illuminates the evolutionary dynamics of this enigmatic clade. Our approach to systematically assess LGT patterns in a broad sense, facilitated by our detection approaches, can be applied to a broad spectrum of genomically characterized eukaryotic clades. Such a comprehensive analysis would deliver a more complete understanding of LGT’s quantitative and qualitative impacts, and would highlight the intricate interplay between biological interactions and genetic exchange in these organisms. Ideally, such a complete picture also builds upon further advances in identifying LGTs, for example also LGTs between closer related lineages, and on the availability of high-quality genome data for a wide range of species. Recent genome sequencing of Ascetosporea, the clade including *M. mackini*, provides a good example thereof, and its observations of primarily gene gains (including from LGT) at recent branches agrees with ours (Supplementary Text, ‘High percentage of LGT in *Mikrocytos mackini*’)^46^. Given ongoing efforts in methodological development in LGT identification^47^ and widely shared calls for broadening genomic sampling of eukaryotes and in particular protists^14,15^, we expect such progress to be on the horizon.

## Methods

### Dataset curation and species phylogeny reconstruction

We analyzed predicted protein sequences from 29 Rhizaria lineages sourced from genomic (n=4) and transcriptomic (n=25) data. *Reticulomyxa filosa* was treated as a transcriptomic dataset for phylogenetic analyses due to its highly fragmented genome assembly.

To infer the Rhizaria species phylogeny, we used the PhyloFisher pipeline^48^ (v.1.0) with default settings. We utilized the ‘phylogenetically-informed’ approach for ortholog search (specifying existing Rhizaria in the dataset as the ‘Blast Seed’) and manually curated orthologs across 238 marker genes using ParaSorter, after excluding H2A and PYGB. The concatenated supermatrix was used to infer a maximum-likelihood species tree in IQ-TREE v.2.0.3^49^ (LG+G4+C60+F model, 1000 ultrafast bootstraps). The tree was arbitrarily rooted on Obazoa+Amoebozoa+CRuMs and Metamonada+Discoba. *Mikrocytos mackini* was manually regrafted as a sister to *Gromia sphaerica* to correct a known long-branch attraction artifact^50^.

### Protein families construction

To mitigate erroneous LGT claims, we additionally compiled a comprehensive background dataset of 418 Stramenopila and Alveolata (SAR) proteomes, representing Rhizaria’s closest relatives (Supplementary Table 1).

**Orthogroup inference:** We used OrthoFinder (v.2.3.8^51^) to group the 418 SAR proteomes, retaining 341,035 orthogroups containing at least one of our 29 focal rhizarian sequences.

**Database downsampling:** To reduce computational load, non-SAR eukaryotic and viral sequences from the NCBI NR database were clustered at 90% identity using CD-hit (v.4.8.1^52^). Bacterial and archaeal sequences from GTDB^53^ (release 89) were downsampled at 70% identity.

**Homology search:** Rhizarian sequences within each orthogroup were used as queries for DIAMOND BLASTP (v2.0.9, ‘ultra-sensitive’ mode, e-value < 0.001). To mitigate taxonomic sampling biases, searches were run separately against seven distinct databases: Archaea, Bacteria, Metazoa, Viridiplantae, Fungi, other eukaryotes, and viruses. A maximum of 2,000 target sequences were retained per search (reduced to 500 for orthogroups exceeding 6,000 members). Rhizarian sequences longer than twice the median length of their orthogroup were excluded to prevent spurious hits from abnormally long sequences.

### Single protein tree inference

Expanded orthogroups containing between 15 and 6,000 sequences were aligned using MAFFT v7.407^54^ (’auto’ mode) and trimmed with BMGE v1.12^55^ (-m BLOSUM30 -b 3 -g 0.7 -h 0.5). Sequences with >80% gaps and alignments with <50 remaining positions were discarded. Maximum-likelihood phylogenies for the remaining 40,951 orthogroups were inferred using IQ-TREE v2.0.3 (LG+F+R5 model) with 1000 SH-aLRT replicates. All resulting trees are available as Newick files in Supplementary Dataset 1.

### Phylogeny-based LGT detection and contamination filtering

Gene trees were parsed using the ETE3 toolkit^56^ to infer the evolutionary origin of rhizarian sequences (vertical, lateral, or invention) via the following step-by-step strict-criteria workflow (Supplementary Figure 8):

**Preprocessing:** Viral and ‘unknown’ sequences were removed. Trees comprising ≥80% rhizarian sequences were classified as rhizarian ‘inventions’.

**Clade definition and merging:** Trees were rooted on a random non-rhizarian leaf. Rhizarian sequences were grouped into monophyletic clades. To account for poor tree resolution, paraphyletic rhizarian clades were merged if separated by <2 strongly supported branches (SH-aLRT ≥ 0.8) or if monophyly was broken by only a single non-rhizarian sequence. Trees with >5 final rhizarian clades were discarded.

**Contamination filtering:** Clades consisting of sequences from a single rhizarian species were subjected to a rigorous contamination check. The entire clade was discarded if any sequence shared >90% identity with any stramenopile/alveolate sequence, or >80% identity with any other taxon (DIAMOND BLASTP: --ultra-sensitive -k 1 --max-hsps 1 --query-cover 70).

**Origin classification:** Internal nodes were taxonomically annotated if they represented the last common ancestor of ≥80% of their descendant leaves (requiring parent branch SH-aLRT > 0.8). The origin was determined by assessing the first and second well-supported parent nodes ascending from the rhizarian clade:

- Vertical origin: Inferred if either parent node was classified as ‘SAR’ or consisted of a diverse range of eukaryotes lacking SAR representation (suggesting gene loss in sister lineages).
- Lateral origin (LGT): Inferred if the descending clades from the parent nodes were exclusively prokaryotic, a specific lower-level non-SAR eukaryotic group (e.g., ‘Metazoa’), or a mix of prokaryotes and eukaryotes.

**Divergence filter:** Rhizarian clades with a median tip-to-tip distance >4 expected substitutions/site to their first sister clade were excluded to prevent erroneous origin inferences from highly divergent sequences..

**Genomic validation:** For *B. natans* and *P. brassicae*, phylogenetically inferred LGTs were only retained if the corresponding gene was located on a genomic scaffold containing at least one gene inferred to have been inherited vertically.

### Machine learning-assisted LGT prediction

**Training data and feature engineering.** To assign evolutionary origins to the substantial fraction of sequences (69%) that were unamenable to phylogenetic inference due to orthogroup size constraints, we developed a sequence-feature-based predictive framework. We first established a ground-truth dataset comprising 35,876 individual sequences whose origins were confidently assigned via our phylogenetic pipeline (21,224 vertical and 14,652 lateral).

Initial benchmarking using a standard HGT index^57^ (the ratio of the DIAMOND bit score of the best ‘alien’ hit to the best ‘native’ hit) yielded only 70% accuracy with a 31% false-positive rate (Supplementary Figure 11A,B), demonstrating the limited utility of sequence similarity scores alone. Consequently, we compiled an array of sequence-intrinsic diagnostic characteristics (e.g., protein length, predicted subcellular localization; Supplementary Table 3) and prepared the data for machine learning by employing one-hot encoding for categorical variables.

**Model selection and hyperparameter tuning.** We evaluated several classifiers from the scikit-learn library^58^ capable of handling mixed categorical and numeric features. A linear discriminant analysis reduced the false-positive rate to 6% but achieved an overall accuracy of only 59% (Supplementary Figure 11C). Ultimately, a Gradient Boosting classifier — an ensemble method utilizing multiple decision trees — emerged as the most effective approach, outperforming other models in accuracy, area under the curve (AUC), and F1 score.

To prevent biases stemming from the disproportionate ratio of vertically inherited to laterally transferred genes, we strictly balanced the two categories within our training dataset. The model was optimized through hyperparameter tuning with five-fold cross-validation on the training set. We finalized the architecture with 1,000 decision trees (n_estimators = 1000) and capped the maximum depth of each tree at 20 to prevent overfitting.

Trained on 80% of the balanced dataset and tested on the remaining 20%, this two-class model (’vertical’ versus ‘lateral’) achieved an overall accuracy of 78% with a 26% false-positive rate (Supplementary Figure 11D). We assessed the influence of individual variables using the feature_importances_ attribute based on Gini impurity (Supplementary Table 3). We also explored a three-class model distinguishing vertical, eukaryotic LGT, and prokaryotic LGT origins, but overall accuracy dropped to 70%, primarily because 27% of vertically inherited genes were misclassified as eukaryotic LGTs (Supplementary Figure 11E) — consistent with the strong structural similarities between these categories (Figure 3B,C). The robust two-class model was therefore retained.

Application and cluster assignment. This optimized classifier was combined with a taxonomic distribution-based prediction to process the unanalyzed orthogroups. Orthogroups comprising >80% rhizarian sequences (excluding viruses) were directly classified as ‘inventions’. The remaining orthogroups were subjected to the Gradient Boosting classifier.

Recognizing that large orthogroups might contain genes from multiple origins, we subdivided their sequences using MCL clustering^59^ with an inflation factor of 1.2. For each resulting cluster, the classifier predicted the origin of each constituent gene and extracted the associated probability. A cluster’s final origin was assigned to the category yielding the highest average probability across its genes, strictly requiring this average to exceed a 70% confidence threshold to discard ambiguous predictions. Finally, to ensure reliability, all predictions were subjected to the identical sequence-identity contamination filters applied in the phylogenetic analysis, including, where possible, verification that the gene resided on a scaffold also containing vertically inherited genes.

### Comprehensive reconstruction of gene evolutionary histories

We reconstructed the evolutionary history of rhizarian genes using two datasets: one from the phylogenetic LGT detection alone (Figure 1, Supplementary Figure 1) and one combining phylogeny-based and ML/distribution-based predictions (Supplementary Figure 2).

For each rhizarian clade (phylogenetic approach) or cluster (prediction approach), we inferred within-Rhizaria gene trees and reconciled them with the species tree. Clades with ≥3 sequences were aligned with MAFFT ‘linsi’, trimmed with BMGE (-m BLOSUM45 -b 3 -g 0.7 -h 0.5), and reconciled using GeneRax v2.0.1^60^ (LG+G model, UndatedDL reconciliation). For two-sequence clades, we used Notung v2.9.1.5^61^; singletons were represented as hypothetical single-branch trees. For prediction-derived clusters, gene trees were inferred with FastTree v2.0^62^ (-lg-gamma) prior to reconciliation. All reconciled trees were annotated with origin type (vertical, lateral, or invention) and, for LGTs, the donor clade.

We then updated each reconciled tree to reflect the full species-tree topology. When a vertically inherited clade rooted on a node more recent than the last common ancestor of Rhizaria (e.g., Foraminifera rather than Rhizaria), we added zero-length branches for the missing ancestral nodes and annotated the corresponding sister lineages as gene losses. This ensured that duplication, transfer, and loss counts were mapped consistently across the complete species tree.

For each node in the updated trees, we recorded: (1) origin (vertical, lateral, invention, or duplication); (2) donor clade, if lateral; (3) median tip distance; and (4) descendant duplication and loss counts. To enable fair comparisons across gene families with different ages and retention patterns, we defined a ‘residence index’: the summed branch length of the species tree pruned to the species retaining that gene. All duplication and loss counts were normalized by this index before aggregation across the species tree.

### Characterizing LGT evolution and function

To compare the evolutionary dynamics and molecular properties of LGT-derived versus vertically inherited genes, we used exclusively the phylogeny-based dataset. For each branch of the species tree, we compared genes whose origin in that specific lineage was lateral against those inherited vertically, excluding genes that entered Rhizaria via LGT in an older ancestor but were subsequently vertically transmitted — as these may have already adapted to the host genome.

**Duplication and loss.** Descendant duplication and loss counts were normalized by the residence index. Losses inferred solely from the ancestral-node reassignment step (above) were excluded. Differences between vertical and lateral genes were assessed with two-sided Mann-Whitney U tests, applied only where both categories contained more than one unique normalized count.

**Sequence divergence.** Root-to-tip median branch lengths were computed for each gene and normalized by the corresponding species-tree root-to-tip distance.

**Domain dynamics.** Pfam domain annotations (HMMER hmmscan against Pfam-A v3.1b2^63^) were obtained for both rhizarian sequences and their sister-clade outgroups. Domain gains and losses were tracked across each within-Rhizaria gene tree and normalized by the residence index.

**Structural and functional annotation.** We predicted a suite of protein attributes for all rhizarian sequences (Supplementary Table 6): intrinsic disorder (MobiDB-lite v3.10.0^64^), coiled-coil regions and protein family assignments (InterProScan v5.48-83.0^65^), subcellular localization (SignalP v5.0b^66^, TargetP v2.0^67^, DeepLoc v1.0^34^), transmembrane domains (Phobius v1.01^68^, TMHMM v2.0c^69^), viral/NCLDV affinities (ViralRecall v2.0^70^), carbohydrate-active enzymes (dbCAN2^71^), and COG categories (eggNOG-mapper v2.1.5^35^). Leaf-level annotations were projected onto internal nodes using Dollo parsimony for qualitative features (requiring presence in ≥10% of leaves) and median values for quantitative features. Annotated features were compared between LGT and vertically inherited genes using chi-square tests (boolean data) or Mann-Whitney U tests (numeric data).

#### Intron content and genomic context

*Introns.* For *B. natans*, *P. brassicae*, and *R. filosa*, we extracted per-gene intron counts, total intron length, intronic proportion, and mean intron length from GFF annotations. For each species, we compared these metrics — and the fraction of genes with ≥1 intron — between vertically inherited and LGT-derived genes, further subdividing the latter by acquisition age (’young’ vs. ‘old’) and donor domain (prokaryotic vs. eukaryotic). Differences were assessed with Mann-Whitney U or chi-square tests.

*Genomic context.* For *B. natans* and *P. brassicae* (*R. filosa* was excluded due to assembly fragmentation), we computed flanking intergenic region (FIR) lengths and distances to the nearest virus-annotated gene (VOG HMM profiles), giant-virus-annotated gene (GVOG HMM profiles), and transposable element on either strand. TEs were annotated with RepeatModeler v2.0.1^72^ (LTR pipeline) and RepeatMasker v4.1.1^73^ (Dfam TE Tools v1.2^74^). Distances were compared between vertical and LGT-derived genes, and among LGT age classes, using Mann-Whitney U tests.

#### Selection and inference of illustrative LGT phylogenies

To select illustrative cases, we identified lineages showing both functional enrichment (COG categories, subcellular localization) and recurrent donor groups among their LGTs. For these candidates, we re-inferred phylogenies under IQ-TREE (LG+C60+G, 1,000 ultrafast bootstraps). For large orthogroups, we retained only the 1,000 non-rhizarian sequences with the shortest branch-length distances to the rhizarian clade in the original tree. After manual validation, we selected two representative cases for Figure 4.

#### Statistics and visualization

All statistical tests were performed with SciPy^75^ (two-tailed where applicable). P-values were corrected for multiple testing using the Benjamini-Hochberg procedure (statsmodels^76^; significance threshold: FDR-adjusted P < 0.05). Figures were generated with ggplot2 and ggtree^77^ in R, except Figure 4, which was produced with iTOL^78^ and rerooted to emphasize the recipient lineage.

## Data availability

Supplementary Figures, Supplementary Tables and Text can be found as Supplementary Material. Supplementary Datasets 1-4 can be found at <u>Figshare</u> (https://figshare.com/projects/Lateral_gene_transfers_LGTs_in_Rhizaria/158240). Figure and table captions, dataset descriptions and Supplementary Text can be found in the file ‘Supplementary Information’.

## Code availability

Scripts for detecting and predicting the origins of Rhizaria genes are available at https://github.com/jolienvanhooff/lgtcallrhizaria

## Acknowledgements

This work was supported by the European Research Council under the European Union’s Horizon 2020 research and innovation programme (ERC Starting Grant to L.E. for Macro-EpiK, 803151). Bioinformatic analyses were run on a local cluster with the help of Philippe Deschamps, on the IFB Core Cluster, the ABiMS Cluster, and National Supercomputer Snellius (SURF Cooperative grant no. EINF-2953). We thank the following people for sharing sequence data of various SAR species: Christian Woehle and Alexandra-Sophie Roy (Kiel University), Rebecca Gast (Woods Hole Oceanographic Institution), Nick Irwin and Varsha Mathur (University of Oxford), Fabien Burki (Uppsala University), Denis Tikhonenkov (Papanin Institute for Biology of Inland Waters), Jürgen Strassert (Leibniz Institute of Freshwater Ecology and Inland Fisheries), Tonje Marita Bjerkan Heggeset (SINTEF Materials and Chemistry), Kristina Terpis and Chris Lane (The University of Rhode Island), and Matthew Brown (Mississippi State University). We thank the PhyloFisher team (Mississippi State University) for providing support in inferring the eukaryotic species phylogeny, Edward Susko (Dalhousie University) for advice on implementing multiple testing correction, and Michael Seidl (Utrecht University) for advice on genomic density estimates. We thank Michelle Leger and Chris Lane for their constructive comments on an early version of this manuscript. We thank David Moreira, Purificación López-García, Andrew Roger, and Courtney Stairs for fruitful discussions.

## Contributions

Conceptualization: J.J.E.v.H. and L.E.; investigation and software development: J.J.E.v.H.; result analysis and interpretation: J.J.E.v.H. and L.E. supervision: L.E.; writing: J.J.E.v.H. and L.E. Funding acquisition: L.E.

## References

1. Ku, C. et al. Endosymbiotic origin and differential loss of eukaryotic genes. Nature 524, 427–432 (2015).

2. Martin, W. F. Too Much Eukaryote LGT. Bioessays 39, 1700115 (2017).

3. Leger, M. M., Eme, L., Stairs, C. W. & Roger, A. J. Demystifying Eukaryote Lateral Gene Transfer (Response to Martin 2017 DOI: 10.1002/bies.201700115). Bioessays (2018) doi:10.1002/bies.201700242.

4. Roger, A. J. Reply to ‘ Eukaryote lateral gene transfer is Lamarckian ‘. Nature Ecology & Evolution 2018 (2018).

5. Van Etten, J. & Bhattacharya, D. Horizontal Gene Transfer in Eukaryotes: Not if, but How Much? Trends Genet. 36, 915–925 (2020).

6. Sibbald, S. J., Eme, L., Archibald, J. M. & Roger, A. J. Lateral Gene Transfer Mechanisms and Pan-genomes in Eukaryotes. Trends Parasitol. (2020) doi:10.1016/j.pt.2020.07.014.

7. Tria, F. D. K. et al. Gene Duplications Trace Mitochondria to the Onset of Eukaryote Complexity. Genome Biol. Evol. 13, (2021).

8. Husnik, F. et al. Horizontal gene transfer from diverse bacteria to an insect genome enables a tripartite nested mealybug symbiosis. Cell 153, 1567–1578 (2013).

9. Shi-Kunne, X., van Kooten, M., Depotter, J. R. L., Thomma, B. P. H. J. & Seidl, M. F. The Genome of the Fungal Pathogen Verticillium dahliae Reveals Extensive Bacterial to Fungal Gene Transfer. Genome Biol. Evol. 11, 855–868 (2019).

10. Eme, L., Gentekaki, E., Curtis, B., Archibald, J. M. & Roger, A. J. Lateral Gene Transfer in the Adaptation of the Anaerobic Parasite Blastocystis to the Gut. Curr. Biol. 27, 807–820 (2017).

11. Xu, F. et al. On the reversibility of parasitism: adaptation to a free-living lifestyle via gene acquisitions in the diplomonad Trepomonas sp. PC1. BMC Biol. 14, 62 (2016).

12. Husnik, F. & McCutcheon, J. P. Functional horizontal gene transfer from bacteria to eukaryotes. Nat. Rev. Microbiol. 16, 67–79 (2018).

13. Sibbald, S. J. & Archibald, J. M. More protist genomes needed. Nature Ecology & Evolution 1, 0145 (2017).

14. Blaxter, M. et al. Why sequence all eukaryotes? Proc. Natl. Acad. Sci. U. S. A. 119, (2022).

15. Schoenle, A. et al. Protist genomics: key to understanding eukaryotic evolution. Trends Genet. 41, 868–882 (2025).

16. Burki, F. & Keeling, P. J. Rhizaria. Curr. Biol. 24, R103–7 (2014).

17. Burki, F. et al. Phylogenomics Reshuffles the Eukaryotic Supergroups. PLoS One 2, e790 (2007).

18. Hackett, J. D. et al. Phylogenomic analysis supports the monophyly of cryptophytes and haptophytes and the association of rhizaria with chromalveolates. Mol. Biol. Evol. 24, 1702–1713 (2007).

19. Burki, F., Roger, A. J., Brown, M. W. & Simpson, A. G. B. The New Tree of Eukaryotes. Trends Ecol. Evol. 35, 43–55 (2020).

20. Strassert, J. F. H., Irisarri, I., Williams, T. A. & Burki, F. A molecular timescale for eukaryote evolution with implications for the origin of red algal-derived plastids. Nat. Commun. 12, 1–13 (2021).

21. Goetz, E. J. et al. Foraminifera as a model of the extensive variability in genome dynamics among eukaryotes. Bioessays 44, e2100267 (2022).

22. Freudenthal, J., Schlegel, M., Bonkowski, M. & Dumack, K. A novel protistan trait database reveals functional redundancy and complementarity in terrestrial protists (Amoebozoa and Rhizaria). Mol. Ecol. Resour. 26, e70064 (2026).

23. Nowack, E. C. M. et al. Gene transfers from diverse bacteria compensate for reductive genome evolution in the chromatophore of Paulinella chromatophora. Proc. Natl. Acad. Sci. U. S. A. 113, 12214–12219 (2016).

24. Vancaester, E., Depuydt, T., Osuna-Cruz, C. M. & Vandepoele, K. Comprehensive and Functional Analysis of Horizontal Gene Transfer Events in Diatoms. Mol. Biol. Evol. 37, 3243–3257 (2020).

25. Shin, N. R., Doucet, D. & Pauchet, Y. Duplication of horizontally acquired GH5_2 enzymes played a central role in the evolution of longhorned beetles. Mol. Biol. Evol. (2022) doi:10.1093/molbev/msac128.

26. Siddique, S. et al. The genome and lifestage-specific transcriptomes of a plant-parasitic nematode and its host reveal susceptibility genes involved in trans-kingdom synthesis of vitamin B5. Nat. Commun. 13, 6190 (2022).

27. Sheikh, S., Fu, C., Brown, M. & Baldauf, S. Deep origins of eukaryotic multicellularity revealed by the Acrasis kona genome and developmental transcriptomes. (2023) doi:10.21203/rs.3.rs-2587723/v1.

28. Sahu, N. et al. Vertical and horizontal gene transfer shaped plant colonization and biomass degradation in the fungal genus Armillaria. Nat Microbiol (2023) doi:10.1038/s41564-023-01448-1.

29. Gilbert, C. & Cordaux, R. Viruses as vectors of horizontal transfer of genetic material in eukaryotes. Curr. Opin. Virol. 25, 16–22 (2017).

30. Irwin, N. A. T., Pittis, A. A., Richards, T. A. & Keeling, P. J. Systematic evaluation of horizontal gene transfer between eukaryotes and viruses. Nat Microbiol (2021) doi:10.1038/s41564-021-01026-3.

31. Fan, X. et al. Phytoplankton pangenome reveals extensive prokaryotic horizontal gene transfer of diverse functions. Science Advances 6, eaba0111 (2020).

32. Ciach, M. A., Pawłowska, J., Górecki, P. & Muszewska, A. The interkingdom horizontal gene transfer in 44 early diverging fungi boosted their metabolic, adaptive, and immune capabilities. Evol. Lett. 8, 526–538 (2024).

33. Nevers, Y., Glover, N. M., Dessimoz, C. & Lecompte, O. Protein length distribution is remarkably uniform across the tree of life. Genome Biol. 24, 135 (2023).

34. Almagro Armenteros, J. J., Sønderby, C. K., Sønderby, S. K., Nielsen, H. & Winther, O. DeepLoc: prediction of protein subcellular localization using deep learning. Bioinformatics 33, 3387–3395 (2017).

35. Cantalapiedra, C. P., Hernández-Plaza, A., Letunic, I., Bork, P. & Huerta-Cepas, J. eggNOG-mapper v2: Functional Annotation, Orthology Assignments, and Domain Prediction at the Metagenomic Scale. Mol. Biol. Evol. (2021) doi:10.1093/molbev/msab293.

36. Nogueira, T. et al. Horizontal gene transfer of the secretome drives the evolution of bacterial cooperation and virulence. Curr. Biol. 19, 1683–1691 (2009).

37. Richards, T. A. & Talbot, N. J. Horizontal gene transfer in osmotrophs: playing with public goods. Nat. Rev. Microbiol. 11, 720–727 (2013).

38. Mertens, J., Aliyu, H. & Cowan, D. A. LEA Proteins and the Evolution of the WHy Domain. Appl. Environ. Microbiol. 84, (2018).

39. Gilbert, C. & Maumus, F. Multiple Horizontal Acquisitions of Plant Genes in the Whitefly Bemisia tabaci. Genome Biol. Evol. 14, (2022).

40. Ropars, J. et al. Adaptive Horizontal Gene Transfers between Multiple Cheese-Associated Fungi. Curr. Biol. 25, 2562–2569 (2015).

41. Habig, M. et al. Frequent horizontal chromosome transfer between asexual fungal insect pathogens. Proc. Natl. Acad. Sci. U. S. A. 121, e2316284121 (2024).

42. Richards, T. A. et al. Horizontal gene transfer facilitated the evolution of plant parasitic mechanisms in the oomycetes. Proceedings of the National Academy of Sciences 108, 15258–15263 (2011).

43. Seczynska, M., Bloor, S., Cuesta, S. M. & Lehner, P. J. Genome surveillance by HUSH-mediated silencing of intronless mobile elements. Nature (2021) doi:10.1038/s41586-021-04228-1.

44. Hibdige, S. G. S., Raimondeau, P., Christin, P.-A. & Dunning, L. T. Widespread lateral gene transfer among grasses. New Phytol. (2021) doi:10.1111/nph.17328.

45. Dunning, L. T. et al. Lateral transfers of large DNA fragments spread functional genes among grasses. Proc. Natl. Acad. Sci. U. S. A. 116, 4416–4425 (2019).

46. Hiltunen Thorén, M., et al. Comparative genomics of Ascetosporea gives new insight into the evolutionary basis for animal parasitism in Rhizaria. BMC Biol. 22, 103 (2024).

47. Morel, B., Williams, T. A., Stamatakis, A. & Szöllősi, G. J. AleRax: a tool for gene and species tree co-estimation and reconciliation under a probabilistic model of gene duplication, transfer, and loss. Bioinformatics 40, btae162 (2024).

48. Tice, A. K. et al. PhyloFisher: A phylogenomic package for resolving eukaryotic relationships. PLoS Biol. 19, e3001365 (2021).

49. Minh, B. Q. et al. IQ-TREE 2: New Models and Efficient Methods for Phylogenetic Inference in the Genomic Era. Mol. Biol. Evol. 37, 1530–1534 (2020).

50. Burki, F. et al. Phylogenomics of the Intracellular Parasite Mikrocytos mackini Reveals Evidence for a Mitosome in Rhizaria. Curr. Biol. 23, 1541–1547 (2013).

51. Emms, D. M. & Kelly, S. OrthoFinder: phylogenetic orthology inference for comparative genomics. Genome Biol. 20, 1–14 (2019).

52. Fu, L., Niu, B., Zhu, Z., Wu, S. & Li, W. CD-HIT: accelerated for clustering the next-generation sequencing data. Bioinformatics 28, 3150–3152 (2012).

53. Parks, D. H. et al. GTDB: an ongoing census of bacterial and archaeal diversity through a phylogenetically consistent, rank normalized and complete genome-based taxonomy. Nucleic Acids Res. 50, D785–D794 (2022).

54. Katoh, K. & Standley, D. M. MAFFT multiple sequence alignment software version 7: improvements in performance and usability. Mol. Biol. Evol. 30, 772–780 (2013).

55. Criscuolo, A. & Gribaldo, S. BMGE (Block Mapping and Gathering with Entropy): a new software for selection of phylogenetic informative regions from multiple sequence alignments. BMC Evol. Biol. 10, 210 (2010).

56. Huerta-Cepas, J., Serra, F. & Bork, P. ETE 3: Reconstruction, Analysis, and Visualization of Phylogenomic Data. Mol. Biol. Evol. 33, 1635–1638 (2016).

57. Boschetti, C. et al. Biochemical diversification through foreign gene expression in bdelloid rotifers. PLoS Genet. 8, e1003035 (2012).

58. Pedregosa, F. et al. Scikit-learn: Machine learning in Python. the Journal of machine Learning research 12, 2825–2830 (2011).

59. Enright, A. J., Van Dongen, S. & Ouzounis, C. A. An efficient algorithm for large-scale detection of protein families. Nucleic Acids Res. 30, 1575–1584 (2002).

60. Morel, B., Kozlov, A. M., Stamatakis, A. & Szöllősi, G. J. GeneRax: A Tool for Species-Tree-Aware Maximum Likelihood-Based Gene Family Tree Inference under Gene Duplication, Transfer, and Loss. Mol. Biol. Evol. 37, 2763–2774 (2020).

61. Stolzer, M. et al. Inferring duplications, losses, transfers and incomplete lineage sorting with nonbinary species trees. Bioinformatics 28, i409–i415 (2012).

62. Price, M. N., Dehal, P. S. & Arkin, A. P. FastTree 2--approximately maximum-likelihood trees for large alignments. PLoS One 5, e9490 (2010).

63. Mistry, J. et al. Pfam: The protein families database in 2021. Nucleic Acids Res. 49, D412–D419 (2021).

64. Necci, M., Piovesan, D., Clementel, D., Dosztányi, Z. & Tosatto, S. C. E. MobiDB-lite 3.0: fast consensus annotation of intrinsic disorder flavours in proteins. Bioinformatics (2020) doi:10.1093/bioinformatics/btaa1045.

65. Jones, P. et al. InterProScan 5: genome-scale protein function classification. Bioinformatics 30, 1236–1240 (2014).

66. Almagro Armenteros, J. J., et al. SignalP 5.0 improves signal peptide predictions using deep neural networks. Nat. Biotechnol. 37, 420–423 (2019).

67. Almagro Armenteros, J. J., et al. Detecting sequence signals in targeting peptides using deep learning. Life Science Alliance 2, e201900429 (2019).

68. Käll, L., Krogh, A. & Sonnhammer, E. L. L. A combined transmembrane topology and signal peptide prediction method. J. Mol. Biol. 338, 1027–1036 (2004).

69. Krogh, A., Larsson, B., von Heijne, G. & Sonnhammer, E. L. L. Predicting transmembrane protein topology with a hidden markov model: application to complete genomes11Edited by F. Cohen. J. Mol. Biol. 305, 567–580 (2001).

70. Aylward, F. O. & Moniruzzaman, M. ViralRecall-A Flexible Command-Line Tool for the Detection of Giant Virus Signatures in ‘Omic Data. Viruses 13, (2021).

71. Zhang, H. et al. dbCAN2: a meta server for automated carbohydrate-active enzyme annotation. Nucleic Acids Res. 46, W95–W101 (2018).

72. Flynn, J. M. et al. RepeatModeler2 for automated genomic discovery of transposable element families. Proc. Natl. Acad. Sci. U. S. A. 117, 9451–9457 (2020).

73. RepeatMasker Home Page. http://www.repeatmasker.org/.

74. TETools: Dfam Transposable Element Tools Docker Container. (Github).

75. Virtanen, P. et al. SciPy 1.0: fundamental algorithms for scientific computing in Python. Nat. Methods 17, 261–272 (2020).

76. Seabold, S. & Perktold, J. Statsmodels: Econometric and statistical modeling with python. in Proceedings of the 9th Python in Science Conference (SciPy, 2010). doi:10.25080/majora-92bf1922-011.

77. Yu, G., Smith, D. K., Zhu, H., Guan, Y. & Lam, T. T.-Y. Ggtree : An r package for visualization and annotation of phylogenetic trees with their covariates and other associated data. Methods Ecol. Evol. 8, 28–36 (2017).

78. Letunic, I. & Bork, P. Interactive Tree Of Life (iTOL) v5: an online tool for phylogenetic tree display and annotation. Nucleic Acids Res. 49, W293–W296 (2021).

