## Supplementary Information for "Eukaryote-to-eukaryote gene transfer pervades the genome evolution of Rhizaria"

### Supplementary Figures

**Supplementary Figure 1. LGT, duplication and invention events across the rhizarian tree of life, excluding single-species inferences.** Similar to Figure 1, except here only proteins present in at least two rhizarian species were analyzed and classified as resulting from vertical inheritance, LGT, or gene invention.

**Supplementary Figure 2. LGT, duplication and invention events across the rhizarian tree of life, combining multiple inference strategies.** Similar to Figure 1 and Supplementary Figure 1, except that it represents results from both the phylogenetics-based strategy to detect the origins of rhizarian proteins, and the machine learning and distribution-based strategy to predict the origins of rhizarian proteins (Methods).

**Supplementary Figure 3**. **Donor frequencies.** For the LGTs presented in Figure 1, the relative frequencies of candidate donors are shown in a pie chart. The donors are classified at the domain level (Eukaryota, Bacteria, and Archaea), or, if such a classification was not possible, as prokaryotes or not determined (ND). n LGT: total number of inferred LGTs at a given branch.

**Supplementary Figure 4. Evolutionary outcomes and genomic contexts of LGTs**. **A.** Percentage of genes with introns in vertically inherited genes compared to LGTs from prokaryotes, combining all LGTs (first red bar from the left) and taxonomically stratified LGTs, representing 'young' (species-specific acquisition, second red bar) to 'old' (acquired by the last common ancestor of Rhizaria, last red bar) acquisitions. Intron presence is depicted for *Bigelowiella natans*, *Plasmodiophora brassicae*, and *Reticulomyxa filosa*. **B.** Distribution of average intron lengths per gene for vertically inherited genes versus LGTs from prokaryotes, combining all LGTs and taxonomically stratified LGTs as in (A). **C.** Rate of sequence evolution in vertically inherited proteins and LGT-derived proteins. For each rhizarian clade in our single gene trees, this sequence evolution was measured as the median branch length from the base of the clade to its tips, and normalized by the median branch length in the species tree. The bars illustrate the mean across proteins. Note that '*Ascetosporea*' received no LGTs (Figure 1), hence this branch lacks data here and in subfigure (D). **D.** Relative numbers of protein domain gain and loss for LGTs versus vertically inherited genes. **E.** Distributions of Flanking Intergenic

Regions (FIRs) associated with LGT genes and vertically inherited genes in *B. natans* and *P. brassicae*. All LGTs are taxonomically stratified as in A,B. **F.** Distributions of distances between predicted transposable elements and LGT genes (all, and taxonomically stratified as in A and B) compared to vertically inherited genes in *B. natans* and *P. brassicae*. **G.** Similar to F, illustrating distances between LGTs or vertical genes to genes predicted to be of giant viruses (NCLDVs) origin. **H.** Similar to F, representing distances to predicted viral sequences of all types. **A, B, E-H:** Clade names for different taxonomic levels (e.g., 'Endomyxa3') are indicated in the phylogeny in Supplementary Figure 3. Asterisks (*) denote statistically significant differences between LGT-acquired genes (red) and vertically inherited genes (blue), determined via the chi-square test of independence (A) or the Mann-Whitney U test (B-H).

**Supplementary Figure 5. Characteristics of genes acquired through LGT compared to vertically inherited proteins. A.** Density plots illustrating coiled-coil lengths of rhizarian proteins encoded by vertically inherited genes and LGT-acquired genes. Mean values for vertically inherited (blue) and LGT (red) genes are indicated by dashed lines. Plots are presented for all LGTs (left panel), LGTs from prokaryotes (middle panel), and from eukaryotes (right panel). P-values were obtained from Mann-Whitney U tests comparing distributions of coiled-coil lengths between LGT and vertically inherited genes. **B.** Similar to A, showing intrinsically disordered region lengths. **C.** Similar to Figure 3B, here combining prokaryotic and eukaryotic LGTs. Comparison of localization propensities between LGT proteins and vertically inherited proteins, indicating the average probability for localization to each organelle as calculated by DeepLoc. A relative localization propensity >1 signifies a stronger on average predicted propensity for LGT proteins to localize to that cellular compartment compared to vertically inherited genes. Predictions were based on annotations of nodes in single gene trees (Methods, 'Characterizing LGT evolution and function using gene evolutionary histories'). P-values were derived from Mann-Whitney U tests comparing distribution probabilities between LGT and vertically inherited proteins. **D.** Similar to Figure 3C, here combining prokaryotic and eukaryotic LGTs. Fold-enrichment of LGT proteins over vertically inherited genes regarding their labeling of a COG functional category based on EggNOG-mapper. Predictions were based on annotations of nodes in single gene trees (Methods, 'Characterizing LGT evolution and function using gene evolutionary histories'). P-values result from chi-square tests of independence comparing the proportion of LGT with a certain annotation and of vertically inherited genes with it. **E-H.** Absolute numbers of LGTs with specific localization and COG functional category predictions from DeepLoc (E, F) and EggNOG-mapper (G, H). Numbers are presented separately for LGTs of putative prokaryotic (E, G) and eukaryotic (F, H) origins. **I.** Similar to Figure 3B, except that it only includes shared (more ancient, not species-specific) LGTs, corresponding to those presented in Supplementary Figure 1. **I,J.** Similar to Figure 3B,C, except that it only includes shared (more ancient, not species-specific) LGTs, corresponding to those presented in Supplementary Figure 1. To enhance visibility and interpretability, we removed the functional category 'Cell motility', because it yielded an extremely high fold enrichment (>10) for eukaryote-derived LGTs, while the absolute values were very small: Only 1 (vertically inherited genes) or 2 (LGT genes) had this annotation and yielded no statistically significant difference. For functional categories 'Nuclear structure' and 'Cytoskeleton', no prokaryote-derived LGT genes were found having these, which is why they lack a bar. In contrast to Figure 3B,C, the here presented values present raw, uncorrected p-values, and therefore, we applied a more stringent cutoff of $P<0.001$.

**Supplementary Figure 6. Relative ages of LGTs.** Distributions of phylogenetic time points ('ancestors') of LGT events that gave rise to the LGT-derived proteins in different lineages, going from the leaf up the species tree, where '0' indicates the time point of the species itself, '1' the shared common ancestor with the nearest sister clade, etc.

**Supplementary Figure 7**. **Eukaryotic tree of life.** This phylogeny was inferred from a concatenated alignment generated with PhyloFisher, using IQ-TREE under the LG+G4+C60+F model of evolution. Branch labels indicate support values estimated from 1000 ultrafast bootstrap replicates. The scale bar indicates the expected number of substitutions per site, where the Mikrocytos mackini branch was manually shrunk to enhance visibility (actual branch length: 3.0). The black-labeled clades and branches were part of the original PhyloFisher dataset (v.1.0), the green-labeled branches represent our examined rhizarians, as presented in Figure 1. The uncollapsed tree can be found on Figshare (https://figshare.com/projects/Lateral_gene_transfers_LGTs_in_Rhizaria/158240, Supplementary Dataset 4). The asterisk signifies that *Euglypha rotunda* was removed from this study (Supplementary Text: "Removal of *Euglypha rotunda* and some *Leptophrys vorax* sequences").

**Supplementary Figure 8**. **Illustration of the gene tree analysis including LGT detection**. "Rhiz": rhizarian. See Methods for details. **A.** Input single gene tree. **B.** Check if >20% of the leaves are comprised of non-rhizarian proteins. If this criterion was not met, we designated this gene as a rhizarian *de novo* invention. **C.** Identify the rhizarian clades in the tree. **D.** Discard single non-rhizarian sequences that split rhizarian clades. **E.** Merge rhizarian clades if fewer than two strongly supported branches (<0.8 SH-like support value) separated them. **F.** Check if there are at most five rhizarian clades in the tree; otherwise the tree was not analyzed. **G.** For rhizarian clades consisting of sequences from a single species, check if the sequences did not have over 80% (for prokaryotes) or 90% (for eukaryotes) sequence similarity to non-rhizarian proteins; otherwise the rhizarian clade was not analyzed further. **H.** Annotate the internal nodes with their taxonomic affiliations at the lowest possible level. **I.** Identify the first two parent nodes of the rhizarian clade having >0.8 branch support. **J.** Determine the origin of the rhizarian clade based on the taxonomic affiliations of the parents (Methods, Supplementary Table 5). **K.** Check if the tip-to-tip distances between rhizarian sequences and their non-rhizarian sisters are smaller than four; otherwise discard the rhizarian clade and its inferred origin. **L.** Validate LGTs in *B. natans* and *P. brassicae* using genome data.

**Supplementary Figure 9. Rhizarian sequence similarity to non-SAR eukaryotic and prokaryotic sequence**. The figure presents DIAMOND blastp percent identity (%) distributions of rhizarian protein sequences compared to two datasets. The top panel shows comparisons against NR excluding SAR (Stramenopila, Alveolata, and Rhizaria) and viral sequences. The bottom panel depicts comparisons against stramenopiles and alveolates from our dataset. Percent identities plotted are those of the best hit in each respective database. The dashed line signifies the cutoffs used for filtering potential contamination (see Methods for details).

**Supplementary Figure 10**. **Workflow outlining the process from initial protein sequence datasets to inferring rhizarian sequence origins, showing data loss along this pipeline.** Green boxes represent the numbers of rhizarian proteins at the beginning and end, while blue boxes depict different protein sets, such as orthogroups or gene trees. Blue arrows between the boxes indicate the applied filtering steps. The small black upward arrow signifies additional data loss: sequences with insufficient alignment coverage (>80% gaps) were removed from gene trees. The '+ non-SAR homologs' black arrow denotes the addition of homologous non-SAR proteins to orthogroups. Black boxes show the percentage of retained proteins at each step. For instance, the 40,952 filtered orthogroups contained 31% of the original rhizarian proteins (835,552). For a detailed description of gene tree analysis (pertinent to the last three blue boxes), including applied filtering steps, see Methods and Supplementary Figure 8.

**Supplementary Figure 11. Examination of HGT index and machine learning methods accuracy in predicting the origins of sequences. A.** Distribution of HGT index among LGT-derived and vertically inherited genes. **B-E.** Accuracies and confusion matrices for different types of classifiers. **B.** HGT index with a cut-off of 1. **C.** linear discriminant classifier based on HGT index. **D.** Two-class gradient boosting classifier based on multiple features. **E.** Three-class gradient boosting classifier based on multiple features. See Methods ('Predicting LGT using protein sequence features') for details.

**Supplementary Figure 12**. **Genetic innovation represented by LGTs**. For each lineage, the fraction of LGT-acquired genes representing completely novel families in rhizarian genomes is represented in yellow; the fraction of LGT-acquired genes having pseudo-paralogs is in blue; the fraction of LGT-acquired genes having pseudo-orthologs is in magenta. See Supplementary Text 'Pseudoparalogy (redundancy), pseudoorthology (displacement)' for details. Between parentheses is the total number of LGTs inferred for each lineage.

### Supplementary Tables

**Supplementary Table 1. Overall genomic and phylogenetic statistics.** The 'SAR dataset' sheet lists the SAR proteomes used here, with reference to the data sources. Other sheets provide absolute and relative counts of proteins analyzed and categorized based on their origins. Sheet 'phylogenetics' is related to Figure 1. It includes counts and percentages of proteins categorized as vertically inherited, laterally acquired through gene transfer (LGT), or originating from gene invention, as determined through phylogenetic analysis. The sheet 'phylogenetics_filter_multispecies' is related to Supplementary Figure 1 and is restricted to rhizarian clades containing sequences from at least two species. The 'phylogenetics_plus_occurrence_machine_learning' sheet is related to Supplementary Figure 2 and combines information from phylogenetic analysis with machine learning or distribution-based prediction (Methods). Vertically inherited proteins are represented in blue columns, LGT-derived proteins in red columns, and invention-derived proteins in yellow columns.

**Supplementary Table 2**. **Rhizarian clades from gene phylogenies, their detected origins and characteristics.** All rhizarian clades resulting from the phylogenetic analysis as presented in Figure 1, with their origins, putative donors and various properties including functional annotations. The annotated phylogenies for rhizarian clades with either an LGT or vertical origin can be found in Supplementary Dataset 2 (NB: these were not generated for rhizarian clades with an ‘invention’, since here the annotation did not affect the origin designation).

**Supplementary Table 3**. **Features used in the gradient boosting classifier for predicting the evolutionary origin of proteins.** The features are ranked based on their importance in the classifier (measured with Gini impurity). Note that the feature set consists of a subset of the features in Supplementary Table 6, supplemented with the HGT index and the last common ancestor ('Recipient lineage) of the represented rhizarian taxa in the rhizarian clade (from the phylogenetic approach, used for training) or cluster (non-phylogenetic, prediction approach, application of the classifier'). The latter feature is split into multiple binary features via one-hot encoding.

**Supplementary Table 4. LGT donors**. Rhizarian lineages with the counts and percentages of LGTs from probable donors at the domain level. Related to Supplementary Figure 3. Lineages were tested for having different ratios of donors relative to the overall distribution (the 'all' row) using the chi-square test, resulting in the listed P-values. The lineages match the ones in the species phylogeny in Supplementary Figure 3.

**Supplementary Table 5**. **LGT detection in gene phylogenies based on parent identities**. Decision schemes for rhizarian clades with one (first sheet) or two (second sheet) valid parents (Methods, Supplementary Figure 8). These schemes embody our rule-based system that, in the phylogenetic detection, determines whether a rhizarian clade is of 'vertical' or 'lateral' (LGT) origin. Note that ‘Group’ here corresponds to the identities acquired in the first annotation step, and can be either prokaryotes, (non-SAR) eukaryotes, SAR, or a mix, whereas ‘Clade’ is based on annotation in a second step, and corresponds to the lowest taxonomic level in NCBI Taxonomy, or GTDB (Methods). The latter is only used to determine if the origin could be an LGT from a particular eukaryotic clade, which is not the case if eukaryotes map to ‘Eukaryota’, i.e. contain sequences from eukaryotic diversity broadly. In the case of the latter, no LGT is inferred, and the origin is ‘vertical’.

**Supplementary Table 6**. **Collections of features to characterize LGTs**. Overview of features used to characterize individual modern-day sequences and, based on that, to project features onto the internal nodes of the gene trees. Two different sets of features were used, i.e. one that is largely qualitative and one that is largely quantitative (detailed in different sheets).

**Supplementary Table 7. Rhizarian clades and clusters from combined analyses, their detected origins and characteristics.** All rhizarian clades and clusters resulting from the phylogenetic analysis and the machine learning- and distribution-based prediction, respectively, with their origins, rhizarian last common ancestor, putative donors, etc. Related to Supplementary Figure 2. Note that the 'inference' column indicates the applied method, either 'second_sister' or 'first_sister', if the phylogenetics-based method was applied, 'rhizaria_dominance', if we applied a prediction based on the occurrence of the protein in Rhizaria only, or 'sequence_classification', if the trained sequence origin classifier was used (see Methods). The column 'prediction_probability' indicates if the average probability for all sequences for a given origin ('vertical' or 'lateral') was larger than the cut-off (set to 0.7, see Methods).

**Supplementary Table 8. Pfam domains fused to LGT-derived proteins**, after acquisition. The LGT-derived sequences were checked for the presence of additional protein domains that were not there yet at the time of the LGT event. The additional domains found to have fused to many LGT proteins can be found at the top of this table, as it is sorted according to the number of such fusion events (see column ‘Number of times this domain was gained by LGT-derived proteins’).

**Supplementary Table 9**. **Protein lengths of LGTs in Rhizaria and donors**. Protein lengths for LGT-derived proteins compared to the entire protein sequences of the “group” (prokaryotes or eukaryotes) to which their putative donor belongs.

| **Protein set 1** | | | | **Protein set 2** | | | | **P-value** |
|---|---|---|---|---|---|---|---|---|
| | **Number** | **Mean AAs** | **Median AAs** | | **Number** | **Mean AAs** | **Median AAs** | |
| All LGT proteins with prokaryotic donor | 9410 | 309 | 229 | Prokaryotes dataset | 32645544 | 308 | 251 | 0.044 |
| All LGT proteins with eukaryotic donor | 15025 | 376 | 280 | Eukaryotes dataset | 25501664 | 434 | 320 | 5.0E-23 |
| *B. natans* LGT proteins with prokaryotic donor | 254 | 442 | 335.5 | Prokaryotes dataset | 32645544 | 308 | 251 | 2.6E-14 |

| | | | | | | | | |
|---|---|---|---|---|---|---|---|---|
| *B. natans* LGT proteins with eukaryotic donor | 502 | 509 | 375 | Eukaryotes dataset | 25501664 | 434 | 320 | 1.2E-08 |
| *P. brassicae* LGT proteins with prokaryotic donor | 89 | 370 | 286 | Prokaryotes dataset | 32645544 | 308 | 251 | 0.028 |
| *P. brassicae* LGT proteins with eukaryotic donor | 276 | 445 | 391 | Eukaryotes dataset | 25501664 | 434 | 320 | 2.6E-04 |

**Supplementary Table 10. Correlations between the numbers of different evolutionary events (LGT, duplication, invention, and loss) and branch length in the species tree**. Events and branch length were measured and collected for each branch. Note that the loss rates are likely overestimated due to proteome incompleteness as indicated by low BUSCO completeness scores (Supplementary Table 1).

| **Event type 1** | **Event type 2** | **Spearman correlation coefficient** | **Spearman correlation *P*-value** |
|---|---|---|---|
| LGT | duplication | 0.77 | $P<0.001$ |
| LGT | invention | 0.66 | $P<0.001$ |
| LGT | branch length | 0.59 | $P<0.001$ |
| LGT | loss | -0.07 | $P=0.86$ |
| duplication | invention | 0.60 | $P<0.001$ |
| duplication | branch length | 0.46 | $P=0.002$ |
| duplication | loss | -0.06 | $P=0.90$ |
| invention | branch length | 0.46 | $P=0.002$ |
| invention | loss | 0.14 | $P=0.54$ |
| branch length | loss | 0.09 | $P=0.78$ |

### Supplementary Text

#### Removal of *Euglypha rotunda* and some *Leptophrys vorax* sequences

Initially, *Euglypha rotunda*, a testate amoeba related to *Paulinella chromatophora* (Supplementary Figure 7), was included in our analyses. Upon analysis, it was found to harbor a substantial number of lateral gene transfers (LGTs), reaching up to 2,738 according to our detection method. These LGTs constituted approximately 90% of the proteins for which we could determine the origin in this organism. Primarily, these candidate LGTs appeared to be from bacteria, particularly planctomycetes. Despite our efforts to filter potentially contaminating sequences (see Methods, 'LGT detection based on single gene trees'), we harbored suspicions regarding the true nature of these apparent LGTs. Consequently, we opted not to include this species in our study.

Furthermore, we observed that *E. rotunda* shared numerous LGTs with the vampyrellid *Leptophrys vorax*, resulting in a remarkably high number of LGTs tracing back to the cercozoan common ancestor. Upon closer scrutiny of the single gene trees, we found it most plausible that the *L. vorax* dataset might have been contaminated with sequences from *E. rotunda*. This was evident as these sequences often branched together with minimal branch length separating them. To address this concern, we filtered sequences from *L. vorax* that could have potentially resulted from such contamination. We identified these sequences by conducting a DIAMOND blastp search of all *L. vorax* protein sequences against all other rhizarian sequences in our dataset. An *L. vorax* sequence was excluded if its best hit was a sequence from *E. rotunda* and if the sequence identity to this hit exceeded 90%.

#### High percentage of LGT in *Mikrocytos mackini*

In our inventory of LGTs, the intracellular oyster parasite *Mikrocytos mackini* stands out with an exceptionally high proportion of LGT-derived proteins, reaching 64% of the phylogenetically analyzed proteins (Figure 1). However, upon closer examination, the majority of these LGTs appear to be species-specific, as evidenced by a significant decrease in this percentage when considering only cases shared with at least one other species (Supplementary Figure 1, 15%). This pattern is further supported by the age estimates of its LGTs (Supplementary Figure 6).

Remarkably, a large proportion of identified LGTs (188 out of 213, 88%) have putative donors within the lineage of the Pacific oyster *Crassostrea gigas*. Initially, this observation raised concerns about potential contamination from transcripts of the animal host, leading to erroneous LGT calls in *M. mackini*. However, our contamination filter (Methods, 'Detecting LGT with single gene trees') effectively eliminated all *M. mackini* sequences with an identity exceeding 80% against any sequence from the NR database, including those of *Crassostrea gigas*.

Therefore, we believe that some LGTs identified in *M. mackini* genuinely reflect transfer events from the host to the parasite, where the LGT proteins diverged over time, as expected.

#### Combined phylogenetics and machine learning approach estimates 8% of proteins were derived from LGT

Our initial investigation encountered a major limitation due to the relatively low proportion of proteins amenable to scrutiny by a phylogenetic approach. Rigorous filtering steps were necessary to prevent the misinterpretation of data with insufficient phylogenetic signal as false positives. Consequently, our results only encompass 12% of all rhizarian proteins in our dataset (Methods, Supplementary Figure 10).

To address this limitation, we sought to predict the origins of proteins not included in the phylogenetic analysis. We considered common non-phylogenetic approaches, such as the 'HGT index'[1] and gene characteristics[2]. However, we found that the HGT index underestimated LGT cases in our dataset (Supplementary Figure 11A, Methods). Additionally, gene characteristics typically applicable to recent LGTs, such as codon bias, were of limited utility in our context. Features like codon biases, which might be different for transferred genes compared to native genes, are only expected to last shortly, as codon preferences quickly adapt to the new genome. Hence, this would not allow us to predict somewhat older LGTs, which is one of the main objectives of our study.

Nevertheless, we identified specific signatures in proteins derived from LGTs (Results, 'Transferred proteins harbor donor-specific signatures'). Leveraging these signatures alongside the HGT index, we employed Gradient Boosting–a machine learning approach– trained on proteins with high-confidence origins determined through phylogenetic analysis (see Methods).

While we employed this Gradient Boosting classifier to proteins with homologs outside of Rhizaria, for proteins (orthogroups) without such homologs we straightforwardly predicted an origination by *de novo* invention, based on their confined distribution.

By integrating results from both phylogenetic and non-phylogenetic approaches, we estimate that 24,807 LGT events occurred across the entire tree of Rhizaria. These gave rise on average to 8% of the examined proteins (which themselves represent 72% of all proteins) in present-day organisms (Supplementary Figure 2, Supplementary Table 7). Notably, this percentage is lower than the 30% and 20% reported for the phylogenetic approach, primarily due to the inclusion of many Rhizaria-specific gene families, which significantly amplify the proportion of inventions to an average of 57% (range: 31-86%). These gene families were typically excluded from phylogenetic-based detection due to containing fewer than 15 sequences, our lower cutoff for inferring a phylogeny.

Furthermore, it is important to acknowledge that the presented number of LGT events may still underestimate the true count due to the incompleteness of most proteomes. Additionally, the trained classifier likely predominantly predicts LGT genes donated by prokaryotes, as they exhibit greater distinction from vertically inherited genes compared to those donated by eukaryotes (see Results, 'Transferred proteins harbor donor-specific signatures' and Methods, 'Predicting LGT using protein sequence features'), suggesting it may underestimate transfer events from eukaryotes.

The machine learning methodology enabled us to delve into the significance of specific features for determining the origin of a protein. This exploration potentially highlights the critical distinctions between genes of vertical and lateral origins. The HGT index emerged as the most significant indicator, followed closely by other key features such as protein length, the likelihood of nuclear localization, and the presence of a signal peptide (see Supplementary Table 3 for more details). Generally, features on predicted subcellular localization were notably more informative compared to those derived from COG functional categories. One possible explanation could be the relative precision of prediction tools for subcellular localization as opposed to the tools available for other predictions, such as COGs.

#### Pseudoparalogy (redundancy), pseudoorthology (displacement)

LGT does not always introduce a gene entirely new to the recipient lineage. It is possible that the recipient genome already encodes a homologous gene, in which case the introduction of the transferred gene results in ‘pseudoparalogy’[3]. Alternatively, related lineages of the recipient one encode a homolog of the transferred gene, suggesting it might have been present in a common ancestor but was lost in the recipient lineage, possibly due to displacement by the transferred gene; this scenario is referred to as 'pseudoorthology'[3]. We aimed to examine the prevalence of these phenomena in Rhizaria.

For each LGT identified phylogenetically, we searched its gene tree for other rhizarian clades that appeared to be of vertical origin. An LGT was deemed 'unique' to the recipient lineage if no other rhizarian clade was present in the tree. However, if rhizarian clades were present and contained sequences from species also present in the LGT clade, this suggested pseudoparalogy. If they only contained different species, this suggested pseudoorthology. Our analysis, conducted separately for each lineage, revealed that 22% of LGTs potentially result in pseudoparalogy, and 24% have one or more pseudoorthologs (Supplementary Figure 12). These findings indicate that both pseudoparalogy and pseudoorthology are relatively frequent phenomena.

The presence of pseudoparalogs could confer several potential fitness advantages to the organism, for example through functional redundancy, which can be advantageous in fluctuating environmental conditions. They can also provide diversification of function, since pseudoparalogs are generally visibly divergent at the sequence level, possibly leading to subtle differences in their functions, allowing organisms to adapt to new niches or environmental challenges by expanding their biochemical and physiological capabilities. Additionally, just like regular paralogs, an increase in the number of gene copies can lead to an increased production of the gene product (gene dosage effect). This can be beneficial in situations where a higher concentration of a particular protein enhances the organism's fitness, such as enzymes involved in metabolism or stress response. Finally, the introduction of pseudoparalogs can increase the regulatory complexity of gene expression, allowing more nuanced control over when and where genes are activated. This can lead to more fine-tuned responses to environmental stimuli or developmental cues.

Similarly, pseudoorthologs can yield selective advantage as they can facilitate rapid adaptation to new environments or niches by providing genetic material that is already somewhat adapted to similar conditions in the donor organism.

#### Introns in prokaryote-derived LGTs

We analyzed various intron-related features for LGTs originating from prokaryotes in three species with sufficiently high-quality genomic assemblies. Our findings, detailed in the main text, reveal that most genes acquired through prokaryotic LGT possess at least one intron (Figure 2A). However, in *B. natans* and *P. brassicae*, these LGT-derived genes are more often devoid of introns compared to genes inherited vertically. When contrasting with extremophilic red algae (19% intron presence in LGT-derived genes versus 42% in native genes[4]) and early-diverging fungi (40% versus 73% in native genes[5]), the incidence of introns in rhizarian LGT genes is higher (Figure 2A: *B. natans* at 81% vs. 89%, *P. brassicae* at 63% vs. 80%, *R. filosa* at 69% vs. 72% for native genes). These findings align with ratios observed across a broad spectrum of SAR species, including *B. natans* and *R. filosa*[6].

Further analysis focused on whether the relatively 'younger' prokaryote-derived LGTs have fewer introns compared to those integrated earlier in the rhizarian evolutionary timeline. Across the three evaluated species, the most recent, species-specific LGTs exhibit a marginally lower frequency of introns than their older counterparts, although the trend is not strongly pronounced, partly due to the minimal LGT acquisition at certain 'older' taxonomic levels (Supplementary Figure 4A).

In addition to intron presence, we examined the lengths of introns within LGT-derived genes, in order to investigate what type of introns get integrated into LGTs. The intron sizes in *P. brassicae* tend to be slightly larger compared to those in vertically inherited genes, while in *R. filosa*, they are smaller (Supplementary Figure 4B presents the median lengths: *B. natans* - 174.5 bp for vertically inherited vs. 176.6 bp for LGT-acquired, P=0.158; *P. brassicae* - 57 bp vs. 58.8 bp, P=0.027; *R. filosa* - 82 bp vs. 72 bp, P=0.002). Despite these minor differences, the lengths are generally similar, suggesting that the introns acquired through LGT resemble conventional spliceosomal introns, and therefore probably are processed by the host's spliceosome, and not through self-splicing mechanisms.

#### LGT sequence divergence

We analyzed the sequence divergence of LGT-acquired genes in their new host species. The analysis was conducted through the examination of branch lengths within single-gene phylogenies, which allowed us to compare the divergence of LGT genes with that of genes inherited vertically. Our findings revealed that in nearly half of the analyzed lineages (13 out of 27, Supplementary Figure 4C), LGT genes exhibited larger branch lengths, suggesting a greater degree of sequence divergence. Moreover, among sixteen lineages where the differences (in any direction) were statistically significant, LGT genes displayed longer branch lengths in ten cases. This pattern suggests that, in certain lineages, LGT genes may undergo accelerated rates of substitutions as they adapt to the cellular environment of their new hosts. The observed increased substitution rate could also be linked to a higher propensity for gene duplication among LGT genes (Figure 2B), as gene duplicates are known to experience faster sequence evolution[7]. Potentially, dynamic evolutionary trajectories, encompassing accelerated sequence evolution and increased gene duplication, contribute to gene integration and adaptation following lateral gene transfer.

#### Domain gain and loss

Beyond sequence divergence, the evolution of genes also encompasses the dynamic process of protein domain gain and loss. After a gene is laterally transferred, it may acquire new protein domains that enable beneficial interactions within the protein network of the recipient species, or lose domains that are not advantageous in its new environment. We employed predictions of Pfam domain compositions at the internal nodes of single gene phylogenies, including the parents of the ancestral rhizarian clade, to track the history of domain gains and losses. This analysis highlighted domain changes associated with LGT, specifically from the parental node to the ancestral rhizarian clade.

Our findings reveal that, in a majority of the examined lineages, genes acquired via LGT tend to gain more domains than those inherited vertically (41 out of 56 lineages, Supplementary Figure 4D). This trend is also evident in the subset of lineages where the differences are statistically significant, with 12 out of 14 lineages showing significantly more domain gains in LGT-acquired genes. Surprisingly, our analysis also showed that these genes experienced a higher rate of domain losses across all lineages (Supplementary Figure 4D), including those where the losses were statistically significant (all 53 lineages). However, this prevalent pattern of domain loss among LGT genes may be linked to the relatively short protein lengths observed in our rhizarian dataset. Protein domains may be truncated (artificially or not) and thus not annotated in the rhizarian proteins, leading to an apparent loss of domains in the gene trees.

To further understand the functional implications of domain gain post-LGT, we investigated which Pfam domains were frequently acquired after transfer. Our observations indicate that LGT genes commonly gain domains predominantly found in eukaryotes, such as Ankyrin repeats, SAM, and WW domains (Supplementary Table 8). These domains are known to facilitate protein-protein interactions, suggesting that domain fusion may enable the integration of new, LGT-acquired genes into the existing protein interaction network of the recipient. Additionally, calcium-binding EF-hand domains and calcium-dependent carbohydrate-binding C-type Lectin domains were also often acquired, suggesting that transferred genes may develop new capabilities for signal transduction and/or protein regulation (through calcium binding), and for carbohydrate binding. This is consistent with findings from early-diverging fungi, where transferred genes often gained carbohydrate-binding domains[5].

#### LGT localization in the genome: nearby transposable elements and viruses

Previous studies have highlighted that LGT-acquired genes often co-localize with ‘parasitic’ genomic elements like retrotransposons, DNA transposons, and integrated viruses, suggesting both a common regulatory repression and the potential for LGTs to be acquired alongside these mobile genetic elements[8]. Here, we assessed the proximity of LGTs to the nearest upstream and downstream transposable elements (TEs), viral sequences, and giant viral sequences. Our analysis revealed minimal significant differences in proximity compared to native genes (Supplementary Figures 4F-H). However, LGTs were more likely to be annotated with a viral origin compared to vertically inherited genes (3.3% versus 2.0%, $P<0.001$), but they were less frequently associated with giant viral annotations (11.4% versus 12.5%, $P=0.017$). This observation suggests that some viruses identified in our gene trees may not only act as vectors for gene transfer but could also serve as the original source of these genes.

#### LGTs from prokaryotes and eukaryotes differ in coiled-coils and disordered regions

As part of our aim to characterize structural features of LGT proteins, we delved into their intrinsically disordered regions and coiled-coil motifs. We observed that, on average,LGT from prokaryotes possess shorter coiled-coil regions and intrinsically disordered regions compared to their vertically inherited counterparts (coiled-coils: 5.0 *vs* 6.8 amino acids in average, $P<0.001$; disordered regions: 27.2 vs 30.0 amino acids in average, $P<0.001$, Supplementary Figure 5A,B, left panels). In contrast, when focusing exclusively on LGTs derived from eukaryotes, the trend reverses, revealing that these LGTs encode proteins with longer coiled-coils (mean 7.9 amino acids, $P<0.001$) and intrinsically disordered regions (mean 40.9 amino acids, $P<0.001$). This pattern aligns with the known higher prevalence of these structural features in eukaryotic proteins compared to prokaryotic ones.

The findings indicate a distinct divergence between prokaryotic and eukaryotic LGTs in terms of their contribution to the recipient proteomes. Specifically, prokaryotic LGTs tend to donate shorter, more globular proteins, whereas eukaryotic LGTs contribute proteins that are more likely to include extended segments of disorder or coiled-coil structures. This distinction underscores the complexity of LGT and its impact on the structural and functional diversity of the recipient proteomes. It also suggests that proteins acquired through LGT do not diverge through a rapid accumulation of additional segments containing disorder or coiled-coil motifs.

#### Protein lengths across datasets

In the section "Transferred proteins harbor demarcating features," we explore the variance in protein lengths between prokaryote-derived LGTs and those acquired from eukaryotes. We found that proteins originating from prokaryotic LGTs are typically shorter than those inherited vertically, whereas eukaryote-derived LGT proteins tend to be longer (Figure 3A). This discrepancy prompted us to examine if the protein length distributions of these LGTs mirror those of their donor organisms. We found that eukaryote-derived LGT proteins were identified to be shorter compared to their counterparts in the donor dataset (Supplementary Table 9). This observation suggests a possible bias towards the transfer of shorter genes or a post-transfer truncation of these proteins. However, it is important to consider the potential impact of technical artifacts on these findings, particularly because proteins derived from transcriptomic data were significantly shorter than those obtained from genomic data, as evidenced in our SAR dataset (see Methods, ‘Assembling a Rhizaria and sister clades dataset’, Supplementary Table 1, column ‘Median protein length’).

To mitigate the influence of such technical discrepancies, we focused our analysis on LGT proteins from species with high-quality genomic data (*B. natans* and *P. brassicae*). Intriguingly, in this refined comparison, LGT proteins were found to be longer than those of their presumed prokaryotic and eukaryotic donors (Supplementary Table 9). This leads us to conclude that the length of LGT-derived proteins is indeed influenced by the phylogenetic background of the donors, but does not strictly align with the protein lengths observed within donor datasets. This discrepancy could indicate a post-transfer elongation of these genes, potentially through the acquisition of additional protein domains discussed above.

Moreover, this confirms that the apparent shortness of many LGT proteins compared to their counterpart in the donor lineage, is likely a consequence of transcriptome sequencing and assembly, which tend to produce artificially short protein lengths. This underscores the importance of considering methodological factors when interpreting data on LGT protein characteristics.

#### Correlations between different types of evolutionary mechanisms

While lineages differed in their frequencies of LGTs, our findings indicated a significant positive correlation between the number of LGT events and gene duplication (Supplementary Table 10, Spearman's correlation, $r=0.770$, $P<0.001$), between LGT and gene invention (Spearman’s correlation: $r=0.656$, $P<0.001$), and between LGT and branch length (Spearman’s correlation: $r=0.588$, $P<0.001$). This pattern suggests that lineages that acquired a substantial number of foreign genes also underwent significant evolutionary changes through gene duplication and rapid sequence evolution. The pattern aligns with putatively similar correlations at the level of individual proteins (‘LGT sequence divergence’).

### Supplementary Data

Available at https://figshare.com/projects/Lateral_gene_transfers_LGTs_in_Rhizaria/158240

**Supplementary Dataset 1**: unprocessed single gene trees from IQ-TREE (n=40951)

**Supplementary Dataset 2**: annotated single gene trees in pdf format for rhizarian clades for which either an LGT or vertical origin was determined (n=32647). The sequence inputs are also provided. More information on the rhizarian clades can be found in Supplementary Table 2.
**Supplementary Dataset 3**: multiple sequence alignments, unprocessed single gene trees from IQ-TREE, and processed trees for Figure 4.
**Supplementary Dataset 4**: full species phylogeny based on PhyloFisher[9] using PhyloFisher dataset v1.0 plus the here added Rhizaria datasets, the former presented in black, the latter presented in green. Related to Supplementary Figure 7. **Euglypha rotunda*: eliminated from our examination (see Supplementary Text: "Removal of *Euglypha rotunda* and some *Leptophrys vorax* sequences").

### References


1. Boschetti, C. *et al.* Biochemical diversification through foreign gene expression in bdelloid rotifers. *PLoS Genet.* **8**, e1003035 (2012).
2. Ravenhall, M., Škunca, N., Lassalle, F. & Dessimoz, C. Inferring horizontal gene transfer. *PLoS Comput. Biol.* **11**, e1004095 (2015).
3. Koonin, E. V. Orthologs, Paralogs, and Evolutionary Genomics. *Annu. Rev. Genet.* **39**, 309–338 (2005).
4. Rossoni, A. W. *et al.* The genomes of polyextremophilic cyanidiales contain 1% horizontally transferred genes with diverse adaptive functions. *Elife* **8**, e45017 (2019).
5. Ciach, M. A., Pawłowska, J., Górecki, P. & Muszewska, A. The interkingdom horizontal gene transfer in 44 early diverging fungi boosted their metabolic, adaptive, and immune capabilities. *Evol. Lett.* **8**, 526–538 (2024).
6. Fan, X. *et al.* Phytoplankton pangenome reveals extensive prokaryotic horizontal gene transfer of diverse functions. *Science Advances* **6**, eaba0111 (2020).
7. Lynch, M. & Conery, J. S. The Evolutionary Fate and Consequences of Duplicate Genes. *Science* **290**, 1151 (2000).
8. Husnik, F. & McCutcheon, J. P. Functional horizontal gene transfer from bacteria to

eukaryotes. *Nat. Rev. Microbiol.* **16**, 67–79 (2018).

9. Tice, A. K. *et al.* PhyloFisher: A phylogenomic package for resolving eukaryotic relationships. *PLoS Biol.* **19**, e3001365 (2021).
